# Elucidating design principles for engineering cell-derived vesicles to inhibit SARS-CoV-2 infection

**DOI:** 10.1101/2021.12.04.471153

**Authors:** Taylor F. Gunnels, Devin M. Stranford, Roxana E. Mitrut, Neha P. Kamat, Joshua N. Leonard

## Abstract

The ability of pathogens to develop drug resistance is a global health challenge. The SARS-CoV-2 virus presents an urgent need wherein several variants of concern resist neutralization by monoclonal antibody therapies and vaccine-induced sera. Decoy nanoparticles—cell-mimicking particles that bind and inhibit virions—are an emerging class of therapeutics that may overcome such drug resistance challenges. To date, we lack quantitative understanding as to how design features impact performance of these therapeutics. To address this gap, here we perform a systematic, comparative evaluation of various biologically-derived nanoscale vesicles, which may be particularly well-suited to sustained or repeated administration in the clinic due to low toxicity, and investigate their potential to inhibit multiple classes of model SARS-CoV-2 virions. A key finding is that such particles exhibit potent antiviral efficacy across multiple manufacturing methods, vesicle subclasses, and virus-decoy binding affinities. In addition, these cell-mimicking vesicles effectively inhibit model SARS-CoV-2 variants that evade monoclonal antibodies and recombinant protein-based decoy inhibitors. This study provides a foundation of knowledge that may guide the design of decoy nanoparticle inhibitors for SARS-CoV-2 and other viral infections.

---

The Coronavirus-19 (COVID-19) pandemic has killed over 5.2M people globally and dramatically underscored the need for therapeutics for treating infectious disease.^1^ The authorization of effective vaccines successfully curbed the pandemic in some locations, but the inability to sufficiently vaccinate the global population and the evolution of highly transmissive, vaccine sera-evading SARS-CoV-2 variants have sustained the COVID-19 pandemic.^2–4^ The development and approval of monoclonal antibody (mAb) therapies that neutralize virions emerged as a viable treatment early in the pandemic and continues to play an important role in treating severe forms of the disease. Unfortunately, circulating viral strains contain mutations in key glycoprotein residues that have rendered several mAb treatments in development, and 60% of the United States Food and Drug Administration (FDA) authorized mAbs, ineffective.^5–7^ Therapies capable of combating circulating and evolving viral strains are urgently needed to supplement the use of vaccines and current antiviral treatments.

A promising complement to mAb therapies is cell-mimicking “decoy” systems—nanoparticles that display host cell receptors on their surface to mimic a cell and bind pathogens.^8^ In the case of SARS-CoV-2, viral entry into a cell is mediated by the binding of the viral Spike glycoprotein (Spike) to the human protein angiotensin-converting enzyme 2 (ACE2) on the cell surface.^9^ As the first step in the viral replication cycle, the Spike-ACE2 binding interaction represents an attractive therapeutic target. In the context of SARS-CoV-2, decoy nanoparticles may exploit this interaction by presenting ACE2 on the decoy surface to bind Spike and inhibit cellular infection by SARS-CoV-2.^10^ This type of strategy has shown promise in other disease contexts, such as in human immunodeficiency virus (HIV) treatment.^11^ The decoy strategy is particularly attractive because it might be robust to evolutionary escape by pathogens^11^—mutations that reduce decoy-pathogen binding affinity (i.e., potential evolutionary escape routes for the virus) will concomitantly attenuate pathogen-cell binding and thus are likely to decrease viral fitness such that these viral variants are unable to outcompete decoy-susceptible variants. Another potential advantage is that if more infectious variants evolve through increasing the affinity with which the pathogen binds a host receptor, such a variant would be equally or more susceptible to inhibition by decoys.

Although any particle that displays a pathogen’s cognate receptor may serve as a decoy particle, biologically-derived nanoparticles that closely resemble the natural host cell microenvironment (e.g., lipid and membrane composition) are of particular interest.^8^ The most prominent biological nanoparticles are extracellular vesicles (EVs)—nanometer-scale particles released by all cells which mediate intercellular transfer of biomolecules.^12, 13^ EVs have recently been investigated as infectious disease decoys^14–17^ in part because, in contrast to most synthetic vehicles, EVs uniquely exhibit low toxicity and low immunogenicity,^13, 18, 19^ and these properties are likely to be of central importance for particles to be administered via sustained infusions or repeat injections, as is envisioned for decoy applications. Although decoy nanoparticles have yet to be evaluated clinically, it seems likely that this approach would be most beneficial for patients experiencing severe or prolonged infections that are not controlled by either their immune system or available antiviral agents.

The COVID-19 pandemic sparked a flurry of decoy EV research that has dramatically advanced the decoy nanoparticle field. Initial speculation that ACE2-containing EVs might inhibit viral infection^20^ was supported in early studies that demonstrated ACE2-EVs bound the SARS-CoV-2 Spike protein^15^ and were capable of inhibiting SARS-CoV-2 pseudotyped lentivirus transduction *in vitro.*^14^ ACE2-EV viral inhibition was later interrogated as a function of dose, providing the community the first quantitative benchmark of decoy potency.^21^ Subsequent work demonstrated efficacy against replication-competent SARS-CoV-2 *in vitro*^16^ and suggested that ACE2-EVs were safe and effective against pseudotyped virus when delivered intranasally in a rodent model.^17^ Most recently, intravenously administered ACE2-EVs lowered the viral load of authentic SARS-CoV-2 in a mouse model, reduced the levels of pro-inflammatory cytokines in lung tissue, and mitigated lung tissue injury.^22^ These studies validated the fundamental concept that decoy EVs could address this disease and raised a number of interesting questions. However, the diversity of experimental systems and designs employed across these studies makes it difficult to synthesize the results of these efforts to evaluate the relationship between specific EV design choices and efficacy. Moreover, we lack understanding as to how decoy EV performance varies across various emerging viral strains, which is an open question of recognized importance.^22^ Resolving these knowledge gaps could help optimize and deploy decoy EV treatments for SARS-CoV-2, novel variants thereof, and perhaps novel viral infections.

In this study, we systematically evaluate the relationships between design features of decoy EVs and performance characteristics *vis-à-vis* inhibition of a model SARS-CoV-2 lentivirus (**Figure 1**). We compare designs across several candidate vesicle subtypes, and we generate new insights into the role of Spike-ACE2 affinity in influencing decoy efficacy. We also compare decoy EVs to an emerging, distinct class of decoy nanoparticles, termed mechanically-generated nanovesicles (NVs). Finally, we evaluate decoy EV-mediated inhibition in the context of several drug-resistant strains of the SARS-CoV-2 Spike protein. These insights will enable future engineering of decoy nanoparticles and provide mechanistic evidence as to how decoy EVs may serve as evolutionarily robust antiviral agents.

**Figure 1.**
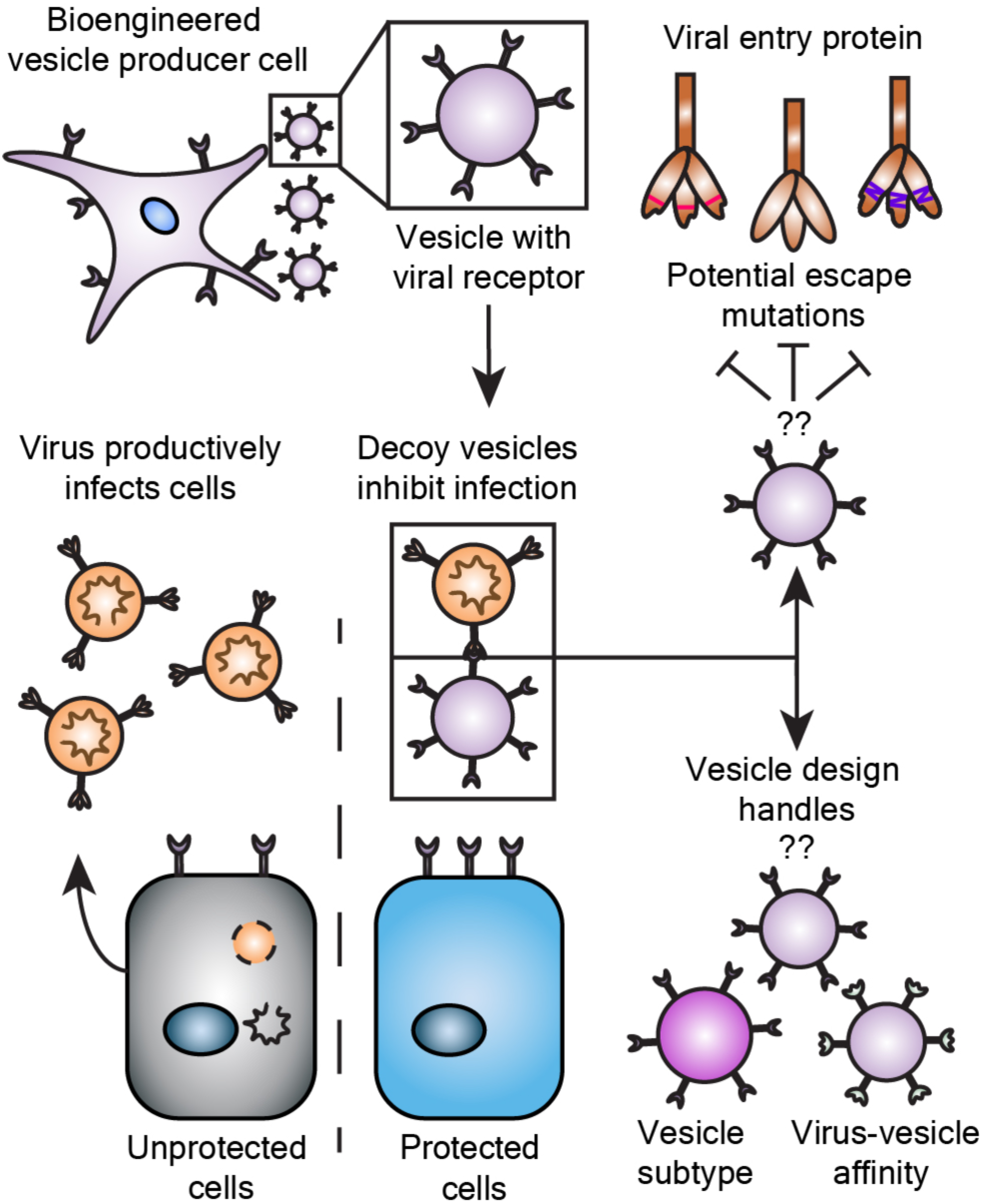
Engineering effective decoy vesicles requires evaluating key design choices. Human cells may be engineered to release vesicles that neutralize virus and inhibit infection. Here, we investigate important open questions as to how general design choices influence the efficacy of decoy vesicle-mediated inhibition of SARS-CoV-2 infection and to what extent this inhibition is robust to mutations that could confer viral escape.

## RESULTS AND DISCUSSION

### Engineered HEK293FT cell lines express high levels of ACE2

To obtain ACE2-containing EVs, we first sought to generate stable cell lines overexpressing ACE2. We engineered HEK293FT cells to stably express a codon-optimized version of the wild-type ACE2 protein (WT-ACE2) via lentiviral-mediated gene delivery. In parallel, we generated a stable cell line expressing a mutant version of the ACE2 gene (Mut-ACE2) that binds to the SARS-CoV-2 Spike protein with higher affinity than does wild-type ACE2 (WT-ACE2) (**Figure S1A**).^23^ Cell lines were analyzed for ACE2 expression, surface display, and EV loading. HEK293FTs did not endogenously express ACE2 at an appreciable level, while both engineered lines expressed high amounts of ACE2 relative to Calu-3s, a model ACE2-expressing lung cell line (**Figure S1B-C**).^14^ Transgenic ACE2 was detected at similar levels across cell lysates from each engineered cell line (**Figure S1C**). We observed a small decrease in apparent molecular weight for the Mut-ACE2 construct relative to WT-ACE2 (**Figure S1C**); this is likely a result of the T92Q mutation which deletes the NXT glycosylation motif at N90.^23^ Surface staining of the cell lines showed high surface expression of ACE2 (**Figure S1D**) which was capable of binding to surface-expressed Spike protein *in trans* (**Figure S2A-B).** We subsequently utilized these engineered HEK293FTs to generate decoy vesicles containing ACE2.

### EVs harvested from engineered HEK293FTs exhibit classical EV characteristics and contain ACE2

Since EVs represent a heterogenous population and various EV subsets can be distinguished by method of purification, we investigated how ACE2 loading varies amongst EV populations. We harvested EVs using differential ultracentrifugation and defined each subset by method of separation, yielding a high-speed centrifugation EV fraction (HS-EVs) and an ultracentrifugation EV fraction (UC-EVs) (**Figure 2A**). Nanoparticle tracking analysis on samples isolated using this protocol revealed two populations of similarly sized nanoparticles (∼100-200 nm), which is a range consistent with reported HEK293FT-dervied EV sizes^24, 25^ (**Figure 2B**). Following established best practices for EV research,^26^ we confirmed that both EV preparations yield particles that exhibit an expected cup-shape morphology by TEM (**Figure 2C**), and both subsets contained standard EV markers CD9, CD81, and Alix **(Figure 2D**). The signal enrichment for CD9 and CD81 blots in UC-EVs versus HS-EVs is consistent with previous reports.^27^ Furthermore, both EV samples were depleted in the endoplasmic reticulum protein calnexin from the producer cells, confirming that our protocol separates cellular debris and EVs.^26^ ACE2 was present in both vesicle populations (**Figure 2E**). We noted that a small ∼18 kDa, C-terminal cleavage product was loaded into EVs along with the full-length protein (**Figure S1B,E**).^28^ Semi-quantitative western blot analysis indicated that, on average, each EV from cells expressing ACE2 (WT or Mut) contained between 500 and 2,500 ACE2 molecules (**Figure S3A-B**). These quantities are comparable to levels reported for expression of engineered proteins on the EV surface.^29^ WT-ACE2 HS-EVs were loaded with more ACE2 than were the rest of the vesicle subtypes (**Figure S3B**), and this effect may be partially attributed to the slightly larger size of HS-EVs compared to UC-EVs (**Figure 2B**). The above characterization steps validated our EV isolation protocol and confirmed the presence of ACE2 in EVs derived from engineered HEK293FTs.

**Figure 2.**
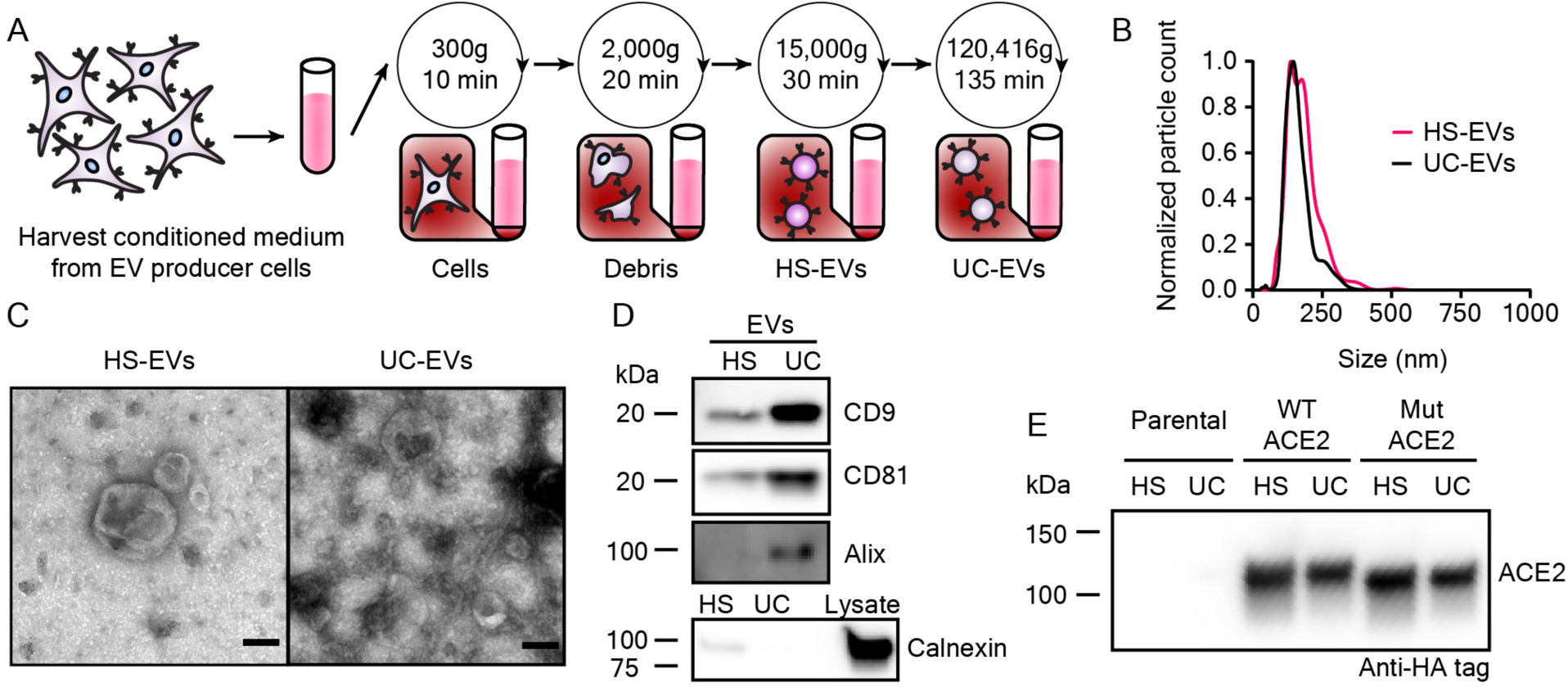
Extracellular vesicles display classical EV characteristics and EVs from engineered cells contain ACE2. A) Depiction of the process used to isolate extracellular vesicles used in this study. B) Representative histogram of nanoparticle tracking analysis of HEK293FT EV subpopulations normalized to the modal value in each population. C) Transmission electron microscopy images of representative EV subpopulations. Scale bar represents 100 nm. D) Western blots of EVs evaluating standard markers CD9, CD81, and Alix, and a blot of EVs and cell lysate evaluting the potentially contaminating endoplasmic reticulum protein, calnexin. E) Western blot against the C-terminal HA-tag of transgenic ACE2 in EV populations from parental or engineered cell lines. Western blots were normalized by vesicle count (for EVs). The Calnexin western blot comparing lysate to EVs used the same number of EVs per well as for CD9, CD81, and Alix and 3 *µ*g cell lysate.

### Modification of virus-producing protocols generates high titer SARS-CoV-2 pseudotyped lentivirus

To evaluate the viral inhibitory potency of decoy EVs, we next developed an *in vitro* transduction assay similar to other reported (**Figure 3A**).^5, 30^ We utilized a 2^nd^ generation lentivirus system to generate lentiviral particles pseudotyped with the SARS-CoV-2 Spike protein (Spike-lenti), wherein an “infection” event causes genomic integration and expression of an enhanced yellow fluorescent protein (EYFP) reporter (i.e., transduction). We chose to use a flag-tagged SARS-CoV-2 Spike construct that contained a D614G mutation and lacked a 19 amino-acid C-terminal sequence because both of these choices have been reported to improve pseudotyping efficiency.^30, 31^ We also generated model recipient cells by engineering HEK293FTs to stably express ACE2. In our hands, this combination produced a detectable but low titer of functional lentivirus: ∼10^1^ transducing units/mL (TU/mL) evaluated on ACE2-expressing HEK293FTs compared to a typical yield of ∼10^5^ TU/mL for a vesicular stomatitis virus G (VSV-G) pseudotyped virus applied to HEK293FT recipient cells (**Figure S4A**). We subsequently explored changing our viral producer cell type from HEK293FTs to HEK293T Lenti-X cells (Lenti-X, Takara), which substantially increased Spike-lenti titer from ∼10^1^ to ∼10^4^ TU/mL (**Figure S4A**). Transduction by Spike-lenti was both Spike and ACE2 dependent **(Figure 3B**). Spike-lenti particles showed no loss of bioactivity after incubation at 37°C for 1 h (**Figure S4B**), which is somewhat surprising given that standard VSV-G pseudotyped lentiviral particles have a reported half-life of ∼30 min when heated in the presence of serum.^32^ ACE2-expressing cells were more susceptible to transduction if cells were plated at the time of transduction rather than the day prior, effectively increasing viral transduction 6-fold (**Figure S4C**). These optimized conditions were used in all subsequent experiments. We anticipate that the procedure reported here to improve viral titer will be useful to the SARS-CoV-2 research community and will circumvent the need for using alternate methods that increase effective viral transduction, such as spinoculation^14^ or adding polybrene,^33^ which could introduce artifacts into the investigation of SARS-CoV-2 infection and its inhibition by decoy nanoparticles.

**Figure 3.**
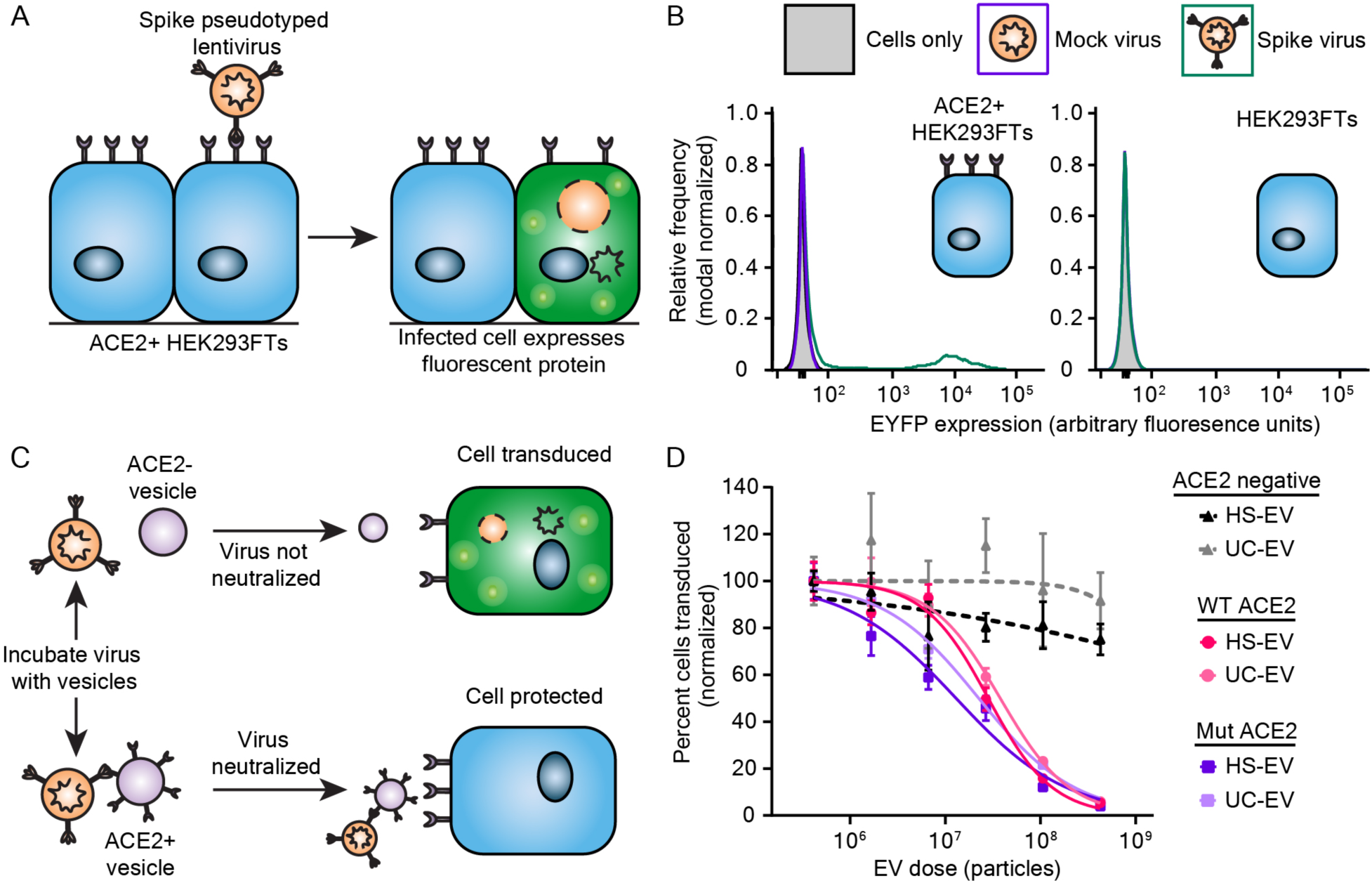
ACE2-containing EVs inhibit pseudotyped SARS-CoV-2 transduction. A) Cartoon depicting pseudotype lentivirus assay. Transduction of ACE2^+^ HEK293FT cells by SARS-CoV-2 Spike pseudotyped virus (Spike-lenti) results in expression of enhanced yellow fluorescent protein (EYFP) as a reporter of viral entry. B) Representative relative frequency histograms of EYFP expression of ACE2^+^ or parental HEK293 FT cells after exposure to no virus, mock virus (no surface glycprotein), or Spike-pseudotyped virus. C) Cartoon depicting pseudotype lentivirus inhibition assay. Effective inhibitors of viral transduction reduce the number of cells expressing EYFP. D) Dose-response curve demonstrating the relationship between EV dose and normalized percentage of cells transduced by Spike-lenti. Curves are normalized to the percent of cells transduced at the lowest EV dose in a particular curve. Symbols repesent the mean of three biological replicates except for the ACE2 negative, HS-EV condition, where the third and fourth dilutions in series are the mean of two biological replicates. Error bars are standard error of the mean. Data are representative of two independent experiments.

### ACE2-containing EVs inhibit Spike-lenti transduction of ACE2-expressing cells in a manner robust to ACE2-Spike affinity and EV subtype

We next quantitatively evaluated the capacity of decoy EVs to inhibit Spike-lenti transduction (**Figure S5)**. Two classes of EVs were prepared (HS-EVs or UC-EVs), either loaded with WT-ACE2 or Mut-ACE2 (**Figure 3C**), yielding four types of ACE2-containing EVs. All four EV types inhibited Spike-lenti transduction in an EV dose-dependent manner. To quantify the potency of these inhibitors, we calculated half-maximal inhibitory dose (ID_50_) values, which were on the order of 1 x 10^7^ particles for all cases (**Figure 3D, S6** and **Table S1, S2**). We note that the ID_50_ metric enables comparisons of EV efficacy within a single assay format (and the same assay format was used throughout this study), but ID_50_ is not an absolute measurement of potency (i.e., this metric is context-specific). Moreover, ID_50_ is a more appropriate metric than is the commonly used IC_50_ metric, since these experiments involve relatively small numbers of discrete particles (i.e., the continuum approximation does not apply). Nonetheless, we analyze both metrics for comparison (**Table S1, S2**). Interestingly, despite detectable differences in ACE2 loading between HS-EVs and UC-EVs (**Figure S3**), potency was similar between HS-EVs and UC-EVs displaying a given ACE2 variant. This finding contradicts a report in which improving ACE2 loading conferred improvements in potency.^22^ Surprisingly, WT-ACE2 and Mut-ACE2 EVs exhibited similar potency despite an estimated 5-10 fold difference in binding affinity of the individual ACE2 variants for Spike,^23^ and this finding held across EV subtypes. One possible interpretation for the similar potency across all four EV-ACE2 combinations is that ACE2 loading for these EV samples is high enough that the effect of loading upon inhibitor potency has saturated (which might not have been the case in prior reports^22^). Control HS-EVs and UC-EVs, which lacked ACE2, had no effect on transduction efficiency, confirming that inhibition of Spike-lenti is ACE2-dependent at all doses evaluated. In addition, EV subtype did not have a meaningful impact on ID_50_, suggesting that vesicle type and purification method are not restrictive design choices for engineering and producing decoy vesicles.

Altogether, our data suggest that decoy EV potency can be relatively independent of intuitively important parameters such as Spike-ACE2 affinity and ACE2 loading levels, but these findings are intimately linked. For example, although the endogenous receptor-virus affinity is ∼20-50 nM for WT-ACE2 and Spike^34, 35^, our particles are loaded with > 500 ACE2 proteins per EV (**Figure S3A-B**), such that the effective avidity of the Spike-lenti-EV interaction is much tighter. These findings should be interpreted in the context of our EV production strategy, where we used a codon-optimized ACE2 gene from a strong promoter (CMV) that is well-suited to the host cell. Altogether these analyses also suggest that it might be possible to generate high potency decoy EVs via a variety of approaches provided that the decoy protein is loaded in sufficient quantity.

### Mechanically generated membrane nanovesicles display similar physical characteristics to EVs and inhibit Spike-lenti in an ACE2-dependent manner

Given the design principles elucidated thus far, we investigated whether other decoy particles could be generated with similar potency using alternative manufacturing approaches, such as lysing ACE2-expressing cells to create membrane nanovesicles (NVs).^10, 36^ One reported benefit of using NVs rather than EVs is that NV manufacturing may yield more vesicles per producer cell in a shorter amount of time.^37^ NVs were generated from WT-ACE2 HEK293FTs via osmotic lysis, sonication, and differential ultracentrifugation (**Figure 4A**).^37–39^ The final membrane pellet was resuspended in PBS and extruded through 100 nm filters to generate ACE2 NVs. Particles exhibited a similar mean size to EVs, although the NV size distribution was narrower (**Figure 4B**). To generate a baseline of protein markers for comparing NVs to EVs, we performed western blots against standard EV markers as in **Figure 2**. Little to no appreciable CD9, CD81, calnexin or Alix were detectable in NVs (**Figure 4C**). ACE2 loading into NVs was also confirmed (**Figure 4D**). Semi-quantitative western blots demonstrated that NVs contained less ACE2 per vesicle than HS-EVs but similar ACE2 per vesicle to UC-EVs derived from WT-ACE2 expressing cells (**Figure S3A-B)**. NVs also exhibited a general cup-shape morphology by TEM (**Figure 4E**), although several NVs displayed internal structures not observed in EVs, which might suggest the presence of multilamellar vesicles in NVs (**Figure S7A**). In our hands, NVs were faster to generate compared to EVs (2-3 days versus 4-5 days, respectively), but particle yields per engineered producer cell seeded were similar (**Figure S7B**). Because WT-ACE2 and Mut-ACE2 Evs similarly inhibited Spike-lenti in earlier experiments (**Figure 3D**), we chose to evaluate NVs using only WT-ACE2. Interestingly, NV decoy potency was remarkably similar to that observed for decoy EVs (**Figures 4F, S8** and **Table S3, S4**). These findings indicate that potent cell-derived decoy nanoparticles can be generated with a variety of biological vesicle subtypes and manufacturing and isolation methods.

**Figure 4.**
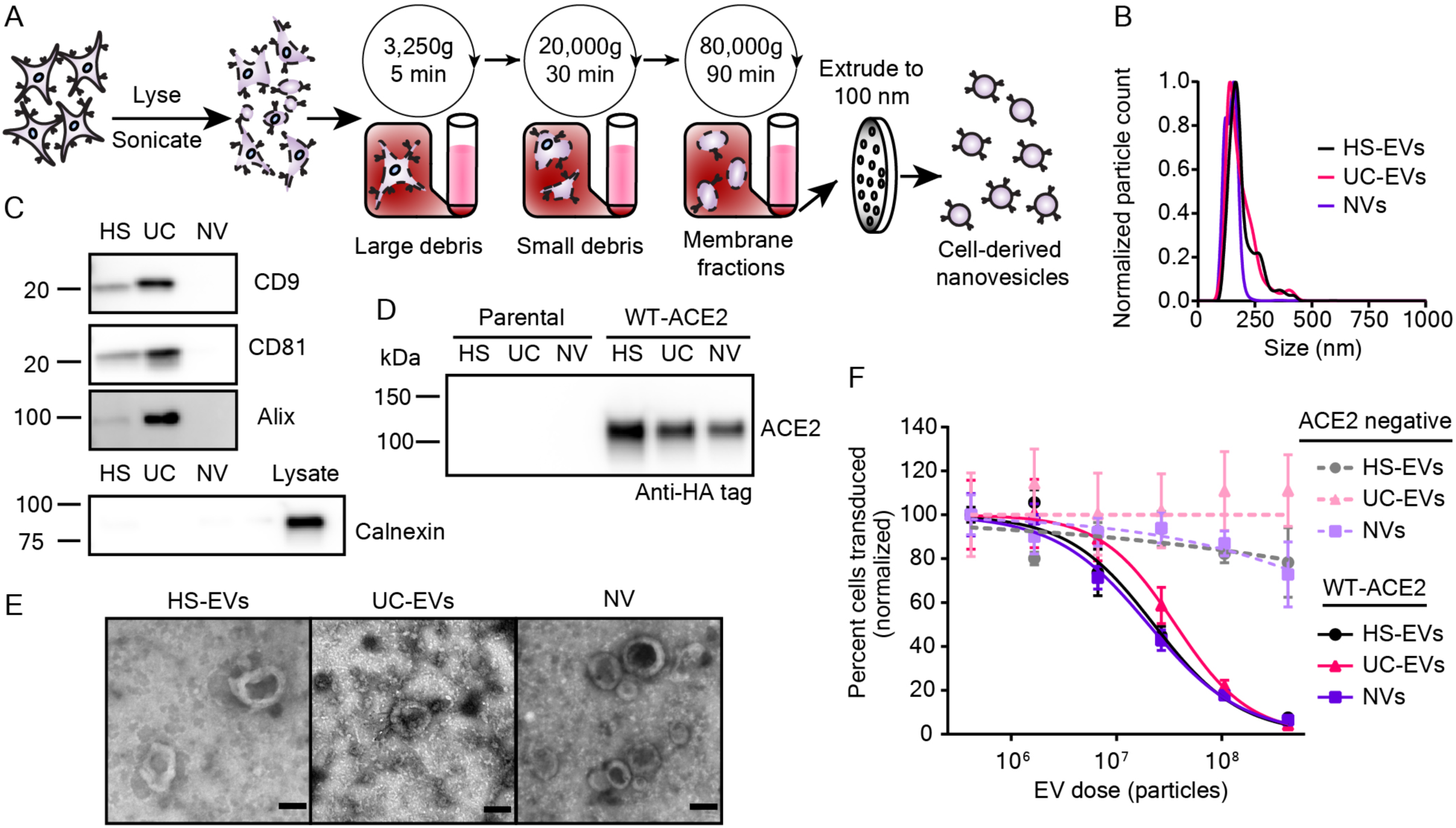
Mechanically generated, cell-derived nanovesicles contain ACE2 and inhibit SARS-Cov-2 pseudotyped lentivirus transduction. A) Cartoon depicting the process used to isolate nanovesicles (NVs) used in this study. B) Representative histogram of nanoparticle tracking analysis of HEK293FT EV subpopulations as compared to NVs; data are normalized to the modal value in each population. C) Western blots of vesicles targeting standard EV markers CD9, CD81, and Alix. Western blot of vesicles and cell lysate of the contaminating endoplasmic reticulum protein, calnexin. D) Western blot against the c-terminal HA-tag of transgenic ACE2 in vesicle populations from parental or engineered cell lines, normalized by vesicle count. Results are representative of two independent experiments. E) Transmission electron microscopy images of NVs alongside micrographs of HS-EVs and UC-EVs. Scale bar represents 100 nm. F) Dose-response curve demonstrating the relationship between decoy vesicle dose and normalized percentage of cells transduced by Spike-lenti. Curves are normalized to the percent of cells transduced at the lowest vesicle dose in a particular curve. Symbols repesent the mean of three biological replicates; error bars are standard error of the mean. Data are representative of two independent experiments.

### ACE2-containing EVs demonstrate broad potency across Spike-lenti variants

Given the diversifying SARS-CoV-2 strains in circulation, we next investigated how decoy EV potency varies across Spike mutant variants. Many such mutants have been identified,^40, 41^ and we decided to first focus our investigation on variants that have demonstrated resistance to drugs that interfere with the Spike-ACE2 interaction, including mAb or sACE2 inhibitors. We identified two Spike mutants that confer such resistance compared to the parental D614G Spike protein (**Figure 5A**). The first mutant contains the SARS-CoV-2 Beta strain’s receptor binding domain (RBD) (Beta), which has three mutations that abolish inhibition by several mAb treatments which are under development or have been authorized by the FDA.^5, 7, 42^ This naturally-evolved viral strain represents a clinically relevant example of a drug-resistant viral strain. The second mutant comprises a point mutation, F486S, that abrogates inhibition by sACE2, likely by altering the Spike RBD and affecting ACE2 receptor engagement (F486S).^43^ Although this mutation has not been reported in circulating SARS-CoV-2 strains, sACE2 treatments currently under development would likely be ineffective against a strain that has or develops this mutation.^44^ We cloned the aforementioned mutants into the same backbone as our D614G (parental) Spike protein and generated Spike-lenti for each variant.

**Figure 5.**
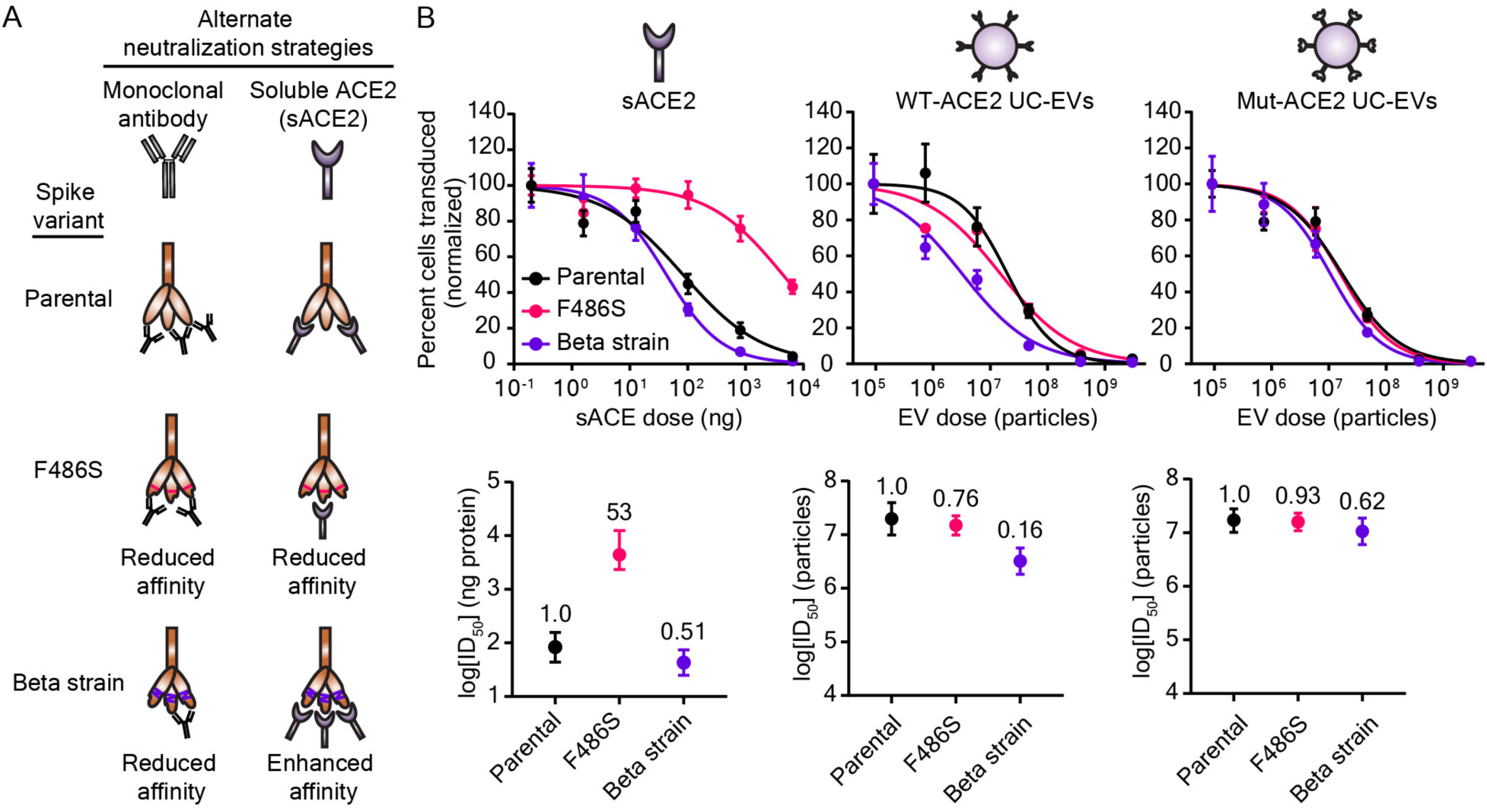
Decoy vesicles are robust to drug-resistant Spike mutations. A) Cartoon depicting the SARS-CoV-2 Spike variants investigated in this study and their relative susceptibility to inhibition by soluble ACE2 (sACE2) or monoclonal antibody (mAb) treatments. B) Top: Dose-response curves depicting the inhibition of various strains of SARS-CoV-2 Spike pseudotyped lentivirus (Spike-lenti) against soluble ACE2, WT-ACE2 UC-EVs, and Mut-ACE2 UC-EVs. Curves are normalized to the percent of cells transduced at the lowest EV dose in a particular curve. Symbols repesent the mean of three biological replicates; error bars are standard error of the mean. Data are representative of two independent experiments. B) Bottom: Log(ID_50_) values calculated from the data in the top portion of this panel. Numbers above each point report the relative resistance for a given strain relative to the parental (D614G) strain for that particular inhibitor treatment. Error bars represent 95% confidence intervals for the parameter (ID_50_) estimation.

We then evaluated the ability of WT-ACE2 UC-EVs, Mut-ACE2 UC-EVs, and sACE to inhibit this panel of Spike-lenti variants (**Figure 5B, S9** and **Table S5-8**). For each treatment, we defined a metric of resistance by dividing the ID_50_ value for the strain considered by the ID_50_ value of a reference, parental strain (in this case, D614G); we term this metric “relative resistance.” A relative resistance value of one indicates that the Spike-lenti variant is equally susceptible to the decoy EV (compared to the reference Spike-lenti), a value less than one indicates that this variant is less resistant to the decoy EV, and a value greater than one indicates that this variant is more resistant to the decoy EV. Relative resistance is defined to facilitate quantitative comparison across strains and treatments by minimizing dependency on virus-to-virus variability (e.g., slight titer differences between samples of virus with different mutations) which can impact ID_50_s. Relative resistance is also useful because it enables comparison across treatments which might be defined in distinct natural units of concentration (e.g., number of vesicles versus mass of sACE2). To evaluate whether any differences in decoy potency were due to different viral titers across viral strains, we also calculated ID_50_s normalized to the viral quantity added (ID_50_/TU) and a relative resistance calculated from ID_50_/TU metrics (relative resistance-TU normalized) in **Table S5 and S7.** Following expected patterns, sACE2 was equally effective against Beta relative to the parental strain, while sACE2 was ∼50 fold less effective against sACE2-resistant F486S Spike-lenti **(Figure 5B**). A particularly promising finding that contrasts with the sACE2 analysis is that all Spike variants analyzed exhibited a relative resistance below 1 for both decoy EV treatments (WT- ACE2 or Mut-ACE2 EVs). Thus, decoy EV potency was robust to Spike mutations known to disrupt the Spike-ACE2 interaction. These data provide the first evidence in a SARS-CoV-2 context that decoy vesicles inhibit viral mutants that are resistant to clinically authorized mAb treatments. Furthermore, these data provide evidence that vesicles displaying wild-type host cell receptors are capable of potently inhibiting viral strains that prove refractory to soluble, protein-based therapeutics. Importantly, this approach circumvents the need for separately engineering a Spike-binding protein in a manner that could be immunogenic and problematic, particularly in the context of sustained or repeated administration.

Given the promising activity of WT-ACE2 EVs against our test strains (which are not prevalent globally), we next sought to extend our investigation to naturally emerging and prominent Spike mutants. In particular, the Delta variant of SARS-CoV-2 rapidly became the dominant strain due to high transmissibility and resistance to mAb and vaccine-induced sera neutralization.^2, 3^ A closely related strain, Delta-plus, demonstrates a similarly high resistance to antibody neutralization and exhibits a marked reduction in affinity for ACE2 relative to wild-type Spike (4-8 fold);^2, 45^ this combination of drug resistance properties integrates features of both the F486S and Beta strain previously investigated. The emerging Lambda variant is less well-studied, but preprints suggest high transmissibility and moderate immune evasion.^46, 47^ Spike-lenti containing the RBD mutations for these variants was generated as follows: Delta (L452R, T478K), Delta-plus: (K417N, L452R, T478K), and Lambda (L452Q, F490S). Viral inhibition experiments were performed with WT-ACE2 UC-EVs using the parental D614G strain as the reference strain (**Figure 6, Table S9**). Notably, the relative resistances of these emerging strains were all below 1, indicating that WT-ACE2 EVs are potent inhibitors of both current and emerging SARS-CoV-2 strains.

**Figure 6.**
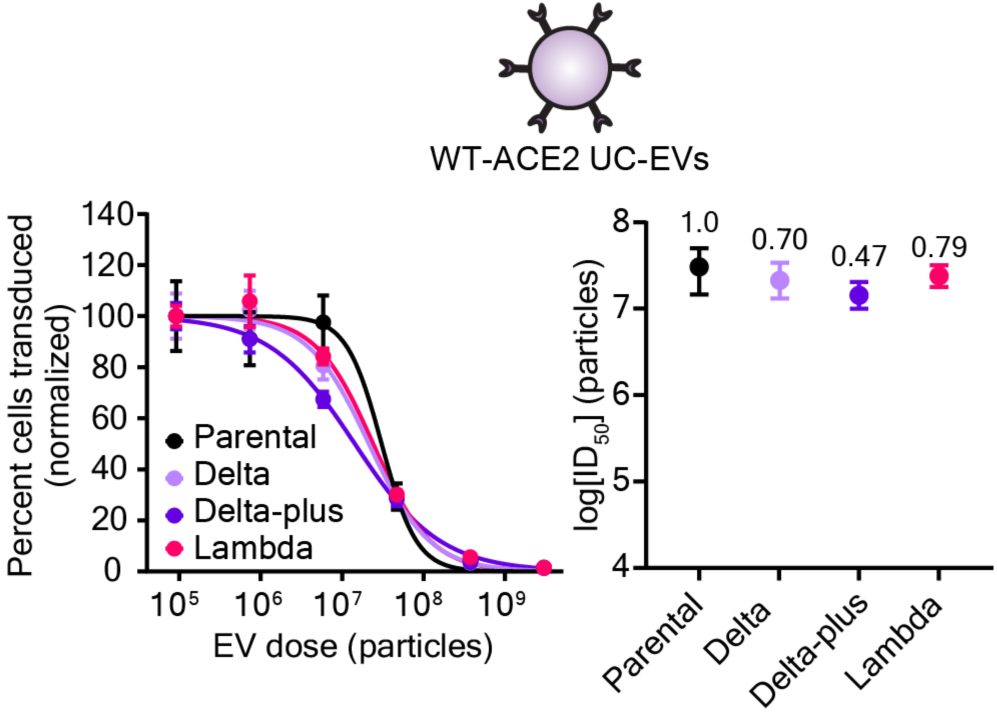
Decoy vesicles confer widespread inhibition of emerging SARS-CoV-2 Spike variants. Left: Dose-response curves depicting the inhibition of various strains of SARS-CoV-2 Spike pseudotyped lentivirus (Spike-Lenti) by WT-ACE2 UC-EVs. The parental strain is the D614G Spike-lenti. Curves are normalized to the percent of cells transduced at the lowest EV dose in a particular curve. Symbols repesent the mean of three biological replicates; error bars are standard error of the mean. Right) Log(ID_50_) values calculated from the dose response curves. Numbers above each point report the relative resistance for a given strain relative to the parental strain (D614G). Error bars represent 99% confidence intervals for the parameter (ID_50_) estimation.

We speculate that avidity is largely responsible for the efficacy of decoy nanoparticles and confers advantages in terms of potency and robustness to drug resistance compared to soluble receptor protein decoys. Given our estimated levels of ACE2 loading (**Figure S3A-B**), our decoy particles contain approximately one ACE2 molecule per 200-500 nm^2^ area of exposed outer membrane, on average. This density of ACE2 display could theoretically facilitate the binding of a decoy vesicle to a virion at several attachment points (e.g., multiple Spike-ACE2 interactions per virion). We hypothesize that the lipid bilayer structure of vesicles enables decoy ACE2 receptors to diffuse across the vesicle surface and improve their likelihood of encountering a Spike protein *in trans*, particularly after an initial vesicle-virion contact has occurred. It is interesting to speculate that decoy particles that contain a fluid bilayer membrane may possess advantages—in terms of either avidity or inhibition mechanism—over nanoparticle systems with fixed protein-attachment points,^48^ but this possibility requires further investigation. By comparing ID_50_ values for vesicle-mediated inhibition (**Figures 5B, S9**)—expressed on a per molecule of ACE2 basis—to ID_50_ values for sACE2-mediated inhibition (**Figures 5B, S9**), we indeed observed evidence of avidity increasing the potency of EVs (**Table S6, S8**). For example, considering the Beta variant Spike-lenti, WT-ACE2 UC-EVs were 51-fold more effective than was sACE2 on a per molecule of ACE2 basis, and Mut-ACE2 UC-EVs were 22-fold more effective than sACE2 (**Table S6)**. This trend also holds for the parental viral strain. This benefit of avidity agrees qualitatively with a previous report comparing sACE2 to ACE2 EVs, but that study found even stronger differences—ACE2-containing EVs were found to be 500-1500 more potent than sACE2.^14^ Differences in experimental setup and the definition of key metrics (e.g., potency, ID_50_) likely account for these quantitative differences, highlighting the importance of evaluating any given design choice using apples-to-apples comparisons. The efficacy of high-avidity particles to inhibit SARS-CoV-2 variants mirrors observations with HIV,^11^ suggesting that this phenomenon could indeed be a general advantage for this type of antiviral inhibitor. An interesting structural consideration is that the SARS-CoV-2 Spike protein quaternary structure is a trimer with three receptor binding domains; all of which may be bound to ACE2 at the same time.^49, 50^ Exploring how various potential modes of avidity and molecular rearrangement with the viral and vesicle membranes contribute to the efficacy of decoy vesicles is an exciting avenue for future research.

## CONCLUSIONS

In this study, we demonstrated that decoy nanoparticles, in the form of ACE2-displaying vesicles, potently inhibit model SARS-CoV-2 viruses bearing drug resistant variants of the Spike protein. We also show that multiple variations of vesicle-based decoys are equally effective *in vitro*, independent of the vesicle subtype or the binding affinity between viral glycoprotein and host cell receptor. Together, these findings suggest that effective inhibitory nanoparticles can be developed using only knowledge of the host cell receptor target of a particular virus. This comparative evaluation informs future preclinical evaluations of this promising approach for potentially treating a wide array of infectious diseases.

## METHODS

### General DNA assembly

Plasmids used in this study were generated using standard polymerase chain reaction (PCR) techniques and/or type II and IIs restriction enzyme cloning. Restriction enzymes, Phusion DNA polymerase, T4 DNA Ligase, and Antarctic phosphatase were purchased from NEB. WT-ACE2 and Mut-ACE2 gene fragments were codon optimized and synthesized by Thermo Fisher and cloned into a pGIPZ backbone (Open Biosciences). The Spike protein from pcDNA3.1-SARS2-Spike was a gift from Fang Li (Addgene plasmid # 145032; http://n2t.net/addgene:145032; RRID:Addgene_145032);^34^ this gene was cloned into a modified pcDNA 3.1 backbone (Clontech-Takara) with a beta-globin intron in the 5’ untranslated region for pseudotyping lentivirus. The Tet3G transactivator (pLVX-EF1a-TET3G) and cognate TRE3GV promoter (pLVX-TRE3G) (Takara) were cloned into modified pGIPZ and pLVX (Takara) backbones, respectively. In this context, the Spike protein (Addgene plasmid # 145032) was cloned downstream of the TRE3G promoter. All plasmids used in this study were sequence-verified, and maps will be published with the final manuscript. Chemically competent TOP10 *Escherichia coli* were used for transformation of all plasmids and subsequently grown at 37°C.

### Plasmid preparation

Plasmid DNA used to generate lentivirus, for viral inhibition assays or for cell-line engineering, was prepared using a polyethlene glycol (PEG) precipitation protocol.^51^ DNA purity and concentrations for relevant experiments were measured with a NanoDrop 2000 (Thermo Fisher Scientific).

### Cell lines and cell culture

HEK293FT cells were purchased from Thermo Fisher/Life Technologies. HEK293T Lenti-X cells were purchased from Takara Bio. Calu-3s were purchased from ATCC (# HTB-55). HEK293FTs and engineered HEK293FTs were grown in a base Dulbecco’s modified eagle medium (DMEM) formulation (Gibco 31600-091). Base medium was further supplemented with 3.5 g/L glucose from Sigma (G7021), 3.7 g/L sodium bicarbonate from Fisher Scientific (S233), and 100 U/mL penicillin and 100 μg/mL streptomycin (15140122), 4 mM L-glutamine (25030-081), and 10% fetal bovine serum (FBS) (16140-071) from Gibco. Lenti-X cells were grown in the HEK293FT formulation supplemented with 1 mM sodium pyruvate from Gibco (11360070). Calu-3s were grown in minimum essential medium from Gibco (41500-018) supplemented with 1.5 g/L sodium bicarbonate and the pH was brought to between 7.0-7.4 with HCl. Calu-3 media was further supplemented with 1 mM sodium pyruvate, 10% FBS, and 100 U/mL penicillin and 100 μg/mL streptomycin. In some cases denoted below, HEK293FTs were briefly cultured in phenol-red free DMEM from Millipore Sigma (D2902). This DMEM formulation was supplemented with 4 mg/L pyridoxine-HCl from Millipore Sigma (P6280), 16 mg/L sodium phosphate from Millipore Sigma (S5011), 3.7 g/L sodium bicarbonate, 3.5 g/L glucose, 100 U/mL penicillin and 100 μg/mL streptomycin, 4 mM L-glutamine (25030-081), and 10% FBS. Cells were maintained in a 37°C incubator held at 5% CO_2_. Spike-expressing cells were induced for at least 24 h prior to assays requiring Spike expression with doxycycline at 1 μg/μL. Doxycyline was from Fisher Scientific (BP2653-5) and resuspended in sterile, nuclease-free water prior to use.

### Cell line generation

HEK293FT cells were used to produce lentivirus for stable cell line generation. 5-6×10^6^ HEK293FTs were plated in 10 cm TC-treated plates and allowed to attach for 5-8 h. Cells were then transfected via calcium phosphate method.^51^ Briefly, DNA (3 μg pMD2.G encoding vesicular stomatitis virus G protein (VSV-G), 8 μg psPAX2 packaging vector, and 10 μg of transfer plasmid encoding desired transgene) were diluted with sterile H_2_O and added to 2M CaCl_2_ to achieve a final concentration of 0.3M CaCl_2_. DNA-containing sample was then added dropwise to an equal-volume of 2x HEPES-buffered saline (280 mM NaCl, 0.5 M HEPES, 1.5 mM Na_2_HPO_4_) and pipetted four times to mix. After 3-4 min, the solution was vigorously pipetted eight times and 2 mL of transfection reagent per 10 cm dish was added dropwise to cells. The plates were gently swirled and incubated overnight at 37°C with 5% CO_2_. The medium was replaced the morning after transfection and cells were incubated for an additional 28-30 h. Conditioned medium containing lentivirus was harvested, clarified via centrifugation at 500 *g* for 2 min at 4°C, and purified through a 0.45 um polyethersulfone filter from VWR (28143-505). Lentivirus was further concentrated via ultracentrifugation at 100,420 *g* for 90 min at 4°C in a Beckman Coulter Optima L-80 XP model and using a SW 41 Ti rotor. Lentivirus was stored on ice until use. 10^5^ HEK293FT parental cells were plated for transduction approximately 24 h in advance in a 12 well TC-treated plate. At the time of transduction, media was aspirated and concentrated lentivirus was added; DMEM was used to bring final volume to 1 mL per well. Two days later, drug selection on cells began and continued for at least one week. ACE2-expressing cell lines were selected using 1 μg/mL puromycin from Invivogen (# ant-pr). Inducible Spike-expressing cell lines were generated from HEK293FTs by inoculating cells with two lentiviruses—one delivering the doxycycline-inducible Tet-On 3G transactivator and one delivering the Spike protein downstream of the TRE3G promoter. These concentrated viruses were added at 1:2 volume ratio, respectively. The cell line was selected using the aforementioned timeline but with 1 μg/mL blasticidin S from Gibco (A11139-03) and 2 μg/mL hygromycin B from Millipore Sigma (400053).

### Cell-binding assays

Cells were grown in 10 cm dishes and harvested with a brief trypsin incubation (< 30 s) followed by quenching with phenol red-free DMEM. Cell suspensions were vortexed to break up clumps, counted, and then diluted to 1 ×10^6^ cells/mL. 100 μL of each cell suspension (if two different cell types were incubated) or 200 μL of the cell suspension (for control wells with only one cell type) were then added to 300 μL phenol red-free DMEM in a non-TC-treated 24 well plate such that the final volume was 500 μL. Cells were incubated at 37°C for 15 min and hand-shaken every 5 min. At 15 min, wells were imaged on a Keyence BZ-x800 microscope using BZ Series Application software v01.01.00.17 and using a PlanApo 4X objective with a numerical aperture of 0.2.

### Surface staining

Two days prior to assay, cells were plated into 12 well tissue culture treated plates such that they were 80-95% confluent at time of harvest. Medium was aspirated, cells were harvested with 1 mL cold fluorescence-activated cell sorting (FACS) buffer (PBS pH 7.4, 2 mM EDTA, 0.05% BSA), and then samples were centrifuged at 150 *g* for 5 min at 4°C. After decanting the supernatant, cells were resuspended in 50 μL FACS buffer and blocked with 10 μL of 1 mg/mL IgG (Thermo Fisher, Human IgG Isotype Control, 02-7102, RRID: AB_2532958) for 5 min at 4°C. After blocking, 2.5 μL of 0.2 μg/μL α-ACE2 antibody from R&D Systems (Human ACE-2 Alexa Fluor® 488-conjugated Antibody, FAB9332G-100UG) was added and incubated for 30 min at 4°C. Cells were washed three times by adding 1 mL cold FACS buffer, centrifuging cells at 150 *g* for 5 min at 4°C, and decanting supernatant. Cells were resuspended in 1 drop of FACS buffer prior to analytical flow cytometry.

### EV production and isolation

15 x 10^6^ HEK293FTs were plated in 15-cm tissue-culture treated plates in 18 mL DMEM. The next morning, the media was replaced with 18 mL of HEK293FT DMEM supplemented with 10% EV-depleted FBS (Gibco, A2720801). After 22-28 h, conditioned medium was harvested as previously reported.^52^ Briefly, the supernatant was clarified by sequential centrifuge spins for 10 min at 300 *g* and 20 min at 2,000 *g*. HS-EVs were pelleted by a subsequent centrifugation at 30 min for 15,000 *g* in a Beckman Coulter Avanti J-26XP centrifuge using a J-LITE JLA 16.25 rotor. The supernatant was centrifuged at 120,416 *g* for 135 min in a Beckman Coulter Optima L-80 XP model using a SW 41 Ti rotor to pellet UC-EVs. All centrifugation was performed at 4°C. EVs were resuspended via gentle pipetting in the conditioned cell medium remaining in their respective vessel.

### Nanoparticle tracking analysis (NTA)

Vesicle concentration and size were measured using a NanoSight NS300 (Malvern) running software v3.4 and a 642 nm laser. Vesicles were diluted to between 2 and 10 x 10^8^ particles/mL in phosphate-buffered saline (PBS) before recording data. Samples were infused at an injection rate setting of 30, imaged with a camera level setting of 14, and analyzed at a detection threshold setting of 7. Three 30 s videos were captured for each sample; vesicle concentrations and size histograms were determined from the average values of the three videos.

### Transmission electron microscopy (TEM)

10 μL of purified vesicles was placed onto a carbon-coated copper grid (Electron Microscopy Services, Hatfield, PA, USA) for 10 min before being wicked away with a piece of filter paper. The grid was dipped in PBS twice to remove excess proteins from the media and was allowed to dry for 2 min. Next, 10 μL of a 2 wt% uranyl acetate solution was placed on the grid for 1 min, before again being wicked away with filter paper. The grid was allowed to fully dry for 3 h to overnight at room temperature. Bright-field TEM imaging was performed on a JEOL 1230 TEM. TEM operated at an acceleration voltage of 100 kV. All TEM images were recorded by a Hamamatsu ORCA side-mounted camera or a Gatan 831 bottom-mounted CCD camera, and AMT imaging software.

### Cell lysate generation

To generate cell lysates, HEK293FTs were washed with cold PBS and lysed with ice-cold RIPA (150 mM NaCl, 50 mM Tris-HCl pH 8.0, 1% Triton X-100, 0.5% sodium deoxycholate, 0.1% sodium dodecyl sulfate) supplemented with protease inhibitor (Pierce/Thermo Fisher #A32953). After a 30 min incubation on ice, lysates were centrifuged at 14,000 *g* for 20 min at 4°C. Protein concentration for each sample was evaluated using a BCA assay. Samples were kept on ice until use or frozen at -80°C for long term storage.

### Cell-derived nanovesicle (NV) generation

HEK293FTs were plated in 10 cm dishes and grown for two days until reaching 80-95% confluency. The day of harvesting, the medium was aspirated and the cells were washed in PBS. Cells were briefly trypsinized (∼30 s) before quenching with EV-depleted DMEM. Cells were pelleted via centrifugation at 150 *g* for 5 min at 4°C and then washed once with ice cold PBS under the same conditions. Cells were resuspended in ice cold PBS, counted using a manual hemocytometer, and pelleted at 150 *g* for 5 min at 4°C. Cells were then resuspended in ice cold lysis buffer (20 mM Tris pH 7.5, 10 mM KCl, 2 mM MgCl_2,_ in nuclease free water supplemented with protease inhibitor tablets)^39^ at a concentration of 0.2-1×10^7^ cells/mL buffer;^37^ typical volumes at this stage were 5-15 mL. Lysis continued on ice for at least 30 min. Samples were then sonicated in an ice-cold water bath (Fisher Scientific, #15337402) at medium power. Samples were sonicated for 10 s and allowed to recover on ice for 50 s; this process was repeated a total of six times such that all samples were sonicated for 1 min. Samples were then clarified via successive centrifugation steps at 4°C in Beckman Coulter Avanti J-26XP centrifuge using either a J-LITE JLA 16.25 rotor or a JA-14.5 rotor: 3,250 *g* for 5 min, and 20,000 *g* for 30 min. Subsequent ultracentrifugation at 80,000 *g* for 90 min pelleted membrane fragments.^38^ PBS was completely aspirated and the samples were resuspended in 30-60 μL PBS per ultracentrifuge tube. Samples were then extruded to 100 nm by passing samples seven times through a 100 nm polycarbonate filter (Whatman #800309) installed in an Avanti Mini Extruder. Samples were then concentrated ∼10X in Amicon Ultra-0.5 mL filter using a 10 KDa molecular weight cutoff (Millipore Sigma #UFC5010) per manufacturer instructions; filters were pre-rinsed with PBS immediately prior to use.

### Western blotting

For western blots comparing the protein content of vesicles, equal numbers of vesicles as determined by NTA were prepared and loaded into the gel (generally 10^8^-10^9^ particles). For western blots comparing the protein content of cell lysates, equal amounts of total protein as determined by BCA were prepared and loaded into gels (generally, 1-10 μg protein). For western blots comparing protein in cell lysates to protein in vesicles, a fixed number of vesicles and a fixed amount of cell lysate were loaded into each well: 4.8 x 10^8^ particles and 3 μg protein, respectively. A detailed western blot protocol has been reported and was followed with the subsequent modifications.^51^ In most cases, the following reducing Laemmli composition was used to boil samples (60 mM Tris-HCl pH 6.8, 10% glycerol, 2% sodium dodecyl sulfate, 100 mM dithiothreitol (DTT), and 0.01% bromophenol blue); in some cases, a non-reducing Laemmli composition (without DTT) was used (**Table S10**). After transfer, membranes were blocked while rocking for 1 h at room temperature in 5% milk in TBST (pH: 7.6, 50 mM Tris, 150 mM NaCl, HCl to pH 7.6, 0.1% Tween). Primary antibody was added in 5% milk in TBST, rocking, for 1 h at room temperature and then washed three times with TBST for 5 min each. Secondary antibody in 5% milk in TBST was added at room temperature for 1 h or overnight at 4°C. Membranes were then washed three times with TBST for 5 min each. The membrane was incubated with Clarity Western ECL substrate (Bio-Rad) and imaged on an Azure c280. Specific antibodies, antibody dilution, heating temperature, heating time, and Laemmli composition for each antibody can be found in **Table S10**.

### Western blot quantification

Digital membrane images were analyzed in ImageJ using the analyze gel function.^53^ Band intensities from ImageJ for the sACE2 standards were analzyed in MATLAB (Mathworks, R2021b) as a function of the amount of ACE2 added in number of molecules (assuming a 115 KDa size for sACE2), and a linear regression was performed to generate a calibration curve. The estimated number of ACE2 proteins in vesicle lanes was determined from the band intensities and calibration curve, and an inverse regression was performed to estimate uncertainty associated with the calibration curve. Estimated number of ACE2 proteins per vesicle was then calculated by dividing the estimated ACE2 proteins per lane by the number of vesicles added to that lane. Error was propagated throughout each calculation, and final error associated with the average number of ACE2 molecules per vesicle was determined by adding-in-quadrature the propagated error and the calculated standard error of the means.

### Pseudotype virus production

HEK293FT or Lenti-X HEK293T cells (Lenti-X) were used to produce SARS-CoV-2 pseudotyped lentivirus (Spike-lenti) for optimizing viral production; Lenti-X cells were used to generate Spike-lenti for all viral inhibition experiments. 5-6×10^6^ Lenti-X cells were plated 24 h prior to transfection of viral plasmids unless otherwise stated; 5-6×10^6^ HEK293FTs were plated in 10 cm TC-treated plates and allowed to attach for 5-8 h. Cells were then transfected via calcium phosphate method as discussed above. Here, 3 μg Spike envelope protein, 8 μg psPAX2 packaging vector, and 10 μg of transfer plasmid encoding an enhanced yellow fluorescent protein (EYFP) transgene were used. The plates were incubated overnight at 37°C with 5% CO_2_. The medium was replaced the morning after transfection and cells were incubated for an additional 32 h prior to harvesting unless otherwise stated. In some cases, medium was replaced, cells were incubated for an additional 24 h, and virus was harvested a second time. Conditioned medium containing lentivirus was harvested, clarified via centrifugation at 500 *g* for 2 min at 4°C, and purified through a 0.45 um polyethersulfone filter (VWR #28143-505). Where required, Spike-lenti was concentrated using Amicon Ultra-15 centrifugal filter units with a 100 kDa cutoff (Millipore Sigma #UFC910024). Samples were centrifuged at 4°C in a Beckman Coulter Avanti J-26XP centrifuge using either a J-LITE JLA 16.25 rotor or a JA-14.5 rotor at 5,000g until concentrated approximately 10-50 fold (generally 10-20 min) and stored on ice at 4°C for up to 1 week or at -80°C until use. To determine functional viral titer, unless otherwise stated, virus was diluted in DMEM and pipetted into a 96 well plate, centrifuged at 500g for 1 min at 4°C to remove bubbles, immediately incubated at 37°C for 1 h. WT-ACE2^+^ HEK23FTs were trypsinized briefly, counted, and 4×10^3^ cells were plated on top of the virus such that the final volume was 200 μL. After 16 h, media was aspirated and 200 μL fresh DMEM was added. Cells were harvested for flow cytometry 3 days after inoculation.

### Viral inhibition assays

Vesicles or soluble ACE2 (Sino Biological, 10108-H08H, resuspended in sterile, nuclease-free H_2_O at 0.25 mg/mL) were serially diluted in DMEM. Spike-pseudotyped lentivirus was added to each sample at a projected MOI between 0.02 – 0.15 and mixed with pipetting. 175 μL of the mixed sample were transferred to TC-treated 96 well plates, centrifuged at 500 *g* for 1 min at 4°C to remove bubbles, and incubated at 37°C for 1 hr. WT-ACE2 expressing HEK293FTs were briefly trypsinized (< 1 min), quenched with DMEM, and counted. Cells were diluted in DMEM and added to plates such that 4 x 10^3^ cells were plated per well in 25 μL media resulting in 200 μL media total per well. Approximately 16 h later, the media was replaced with 200 μL fresh DMEM and the cells were cultured for an additional two days (∼72 h post inoculation). Cells were harvested for flow cytometry via trypsin, quenched with phenol red-free DMEM, and diluted with at least 5 volumes of FACS buffer in FACS tubes. Samples were centrifuged at 150 *g* for 5 min at 4°C, the supernatant was decanted, and the samples were stored at 4°C until flow cytometry analysis.

### Flow cytometry and analysis

Analytical flow cytometry was performed on a BD LSR Fortessa Special Order Research Product (Robert H. Lurie Cancer Center Flow Cytometry Core); EYFP expression and Alexa Fluor® 488 staining was measured using the fluorescein isothiocyanate (FITC) channel from a 488 nm excitation laser and captured using a 505 nm long pass filter and a 530/30 nm bandpass filter. Approximately 5,000-10,000 single cells were analyzed for each sample on FlowJo software v10. As illustrated in **Figure S5**, cells were identified using side scatter versus forward scatter gating, and singlets were isolated using forward scatter-height versus forward scatter-area. In transduction experiments, cells without viral treatment were used as a fluorescent gating control such that < 0.5% of these cells were gated as EYFP+. The output metric for each sample in such experiments was percent of single cells that were transduced (EYFP+). In dose-response curve experiments, the percent of transduced cells for a given treatment was normalized by the percent of transduced cells determined from that treatment’s largest dilution as depicted in **Figure S5**.^54^ Curves were then fit with a four parameter, nonlinear regression in GraphPad Prism 9.2. Convergence criteria was set to “Strict” with 10,000 maximum iterations, and the regression was constrained as follows: “Bottom” = 0, “Top” = 100, “IC50” > 0.

## ASSOCIATED CONTENT

### Supporting Information

Characterization of engineered ACE2 expression cells, ACE2-Spike cell binding assay micrographs, semi-quantitative ACE2 western blot analyses on EVs and NVs, pseudotyped lentivirus optimization; data analysis workflow for viral inhibition experiments; characterization of NVs; replicate experiments for Figure 3D, Figure 4F, Figure 5B; tables describing ID_50_, ID_50_/TU, IC_50_, and relative resistance parameters for all viral inhibition experiments; table describing western blot conditions and the antibodies used. File type: PDF.

### Data Availability

Datasets generated and/or analyzed in this study are available from the corresponding authors upon reasonable request. Raw experimental data for all figures will be published with the final manuscript. Plasmid maps and an annotated description of plasmids used in this study will be published with the final manuscript. Key plasmids used in this study will be deposited at Addgene.

## Author Contributions

T.F.G., D.M.S., N.P.K., and J.N.L. conceived the initial project. T.F.G. performed the experiments. R.E.M. performed transmission electron microscopy. T.F.G., D.M.S., N.P.K., and J.N.L. planned and analyzed experiments. T.F.G., D.M.S., N.P.K., and J.N.L. wrote the manuscript. N.P.K., and J.N.L. supervised the work.

## Notes

The authors declare no competing financial interests.

## ACKNOWLEDGMENT

This work was supported in part by the National Science Foundation under Grant No. 1844219 (NK, JNL) and 1844336 (NK) and a gift from Kairos (JNL). This work was supported by the Northwestern University – Flow Cytometry Core Facility supported by Cancer Center Support Grant (NCI CA060553). This work made use of the BioCryo facility of Northwestern University’s NUANCE Center, which has received support from the SHyNE Resource (NSF ECCS-2025633), the IIN, and Northwestern’s MRSEC program (NSF DMR-1720139). This material is based upon work supported by the National Science Foundation Graduate Research Fellowship under Grant No. (T.F.G. # DGE-1842165 and D.M.S. #DGE-1324585). T.F.G. and R.E.M. were supported in part by the Northwestern University Graduate School Cluster in Biotechnology, Systems, and Synthetic Biology, which is affiliated with the Biotechnology Training Program. REM was supported in part by the National Institutes of Health Training Grant (T32GM008449) through Northwestern University’s Biotechnology Training Program. Biological and chemical analysis] was performed in the Analytical bioNanoTechnology Core Facility of the Simpson Querrey Institute at Northwestern University. The U.S. Army Research Office, the U.S. Army Medical Research and Materiel Command, and Northwestern University provided funding to develop this facility and ongoing support is being received from the Soft and Hybrid Nanotechnology Experimental (SHyNE) Resource (NSF ECCS-1542205). This work was supported by the Northwestern University Sanger Sequencing Facility. Any opinion, findings, and conclusions or recommendations expressed in this material are those of the authors(s) and do not necessarily reflect the views of the National Science Foundation. The authors kindly thank the Kamat and Leonard lab members for useful discussions throughout the planning, experimental, analysis, and writing phases of this project.

## Notes

### Competing Interest Statement

The authors have declared no competing interest.

### Summary of Updates

Corrected bioRxiv listing (not manuscript) to include a missing author.

