## Supporting Information for "Elucidating design principles for engineering cell-derived vesicles to inhibit SARS-CoV-2 infection"

for

**Contents:**

- Supplementary Figures 1-10
- Supplementary Tables 1-10

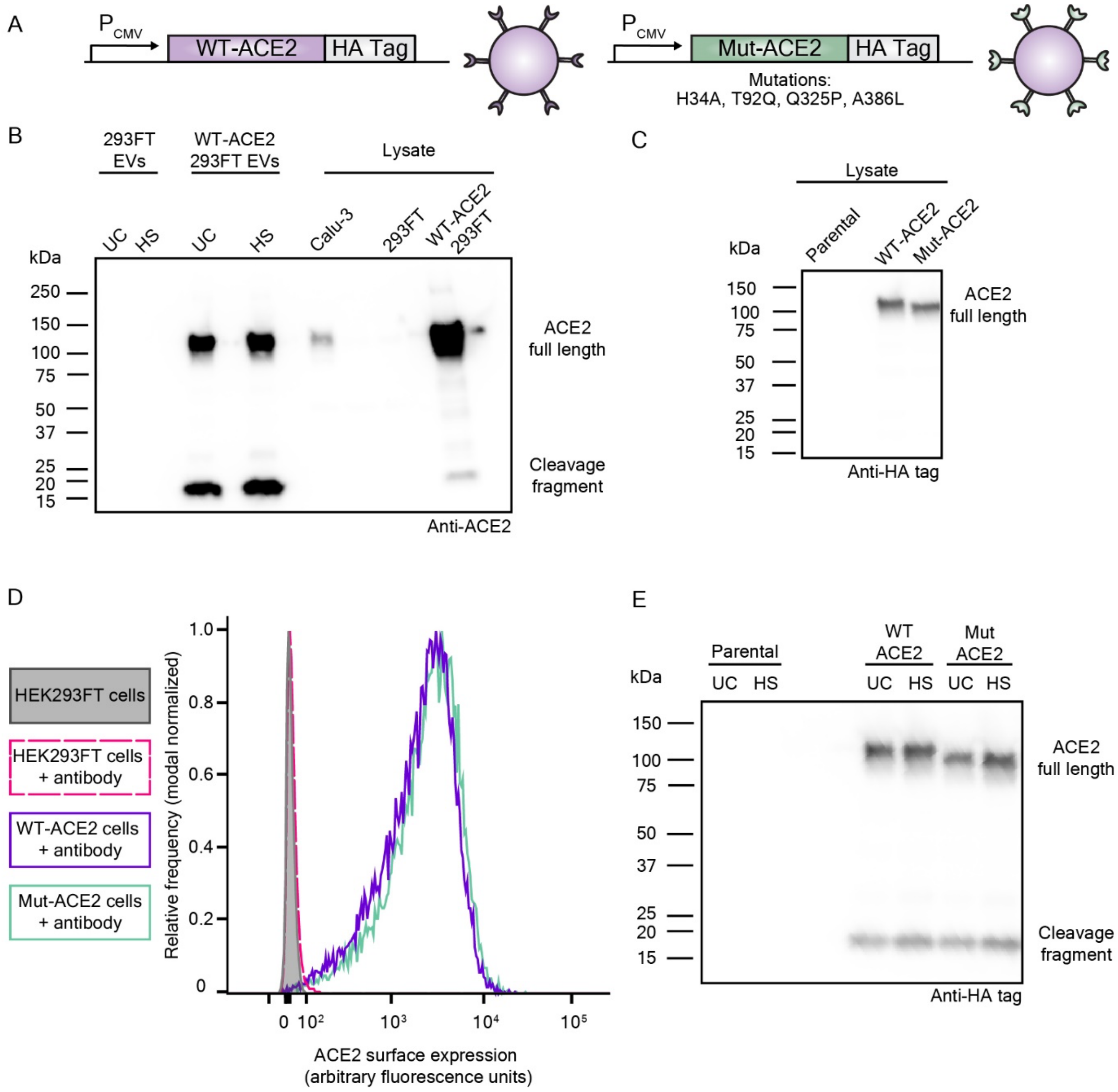


**Supplementary Figure 1.** Engineering and characterization of ACE2-expressing cells and ACE2-containing EVs. A) Schematic depicting the ACE2 DNA constructs used in the study and a cartoon of how the constructs are displayed on vesicles. Both WT-ACE2 and Mut-ACE2 are codon optimized, downstream of a CMV promoter, and fused to a C-terminal HA tag. Mut-ACE2's specific amino acid mutations are identified below the DNA cartoon. B) Western blot using an α-ACE2 primary antibody to detect ACE2 in EVs and cell lysates in non-engineered HEK293FTs and engineered HEK293FTs. Calu-3s are known to express ACE2. C) Western blot using an α-

HA-tag primary antibody to detect transgenic ACE2 in engineered cell lysates. D) Flow cytometric analysis of surface staining of parental or engineered cells with a fluorescent, α-ACE2 primary antibody. E) Western blot using an α-HA-tag primary antibody to detect transgenic ACE2 in engineered EVs. Within each western blot, equal numbers of particles (for EVs) or amount of protein (for lysate) were loaded in each well.

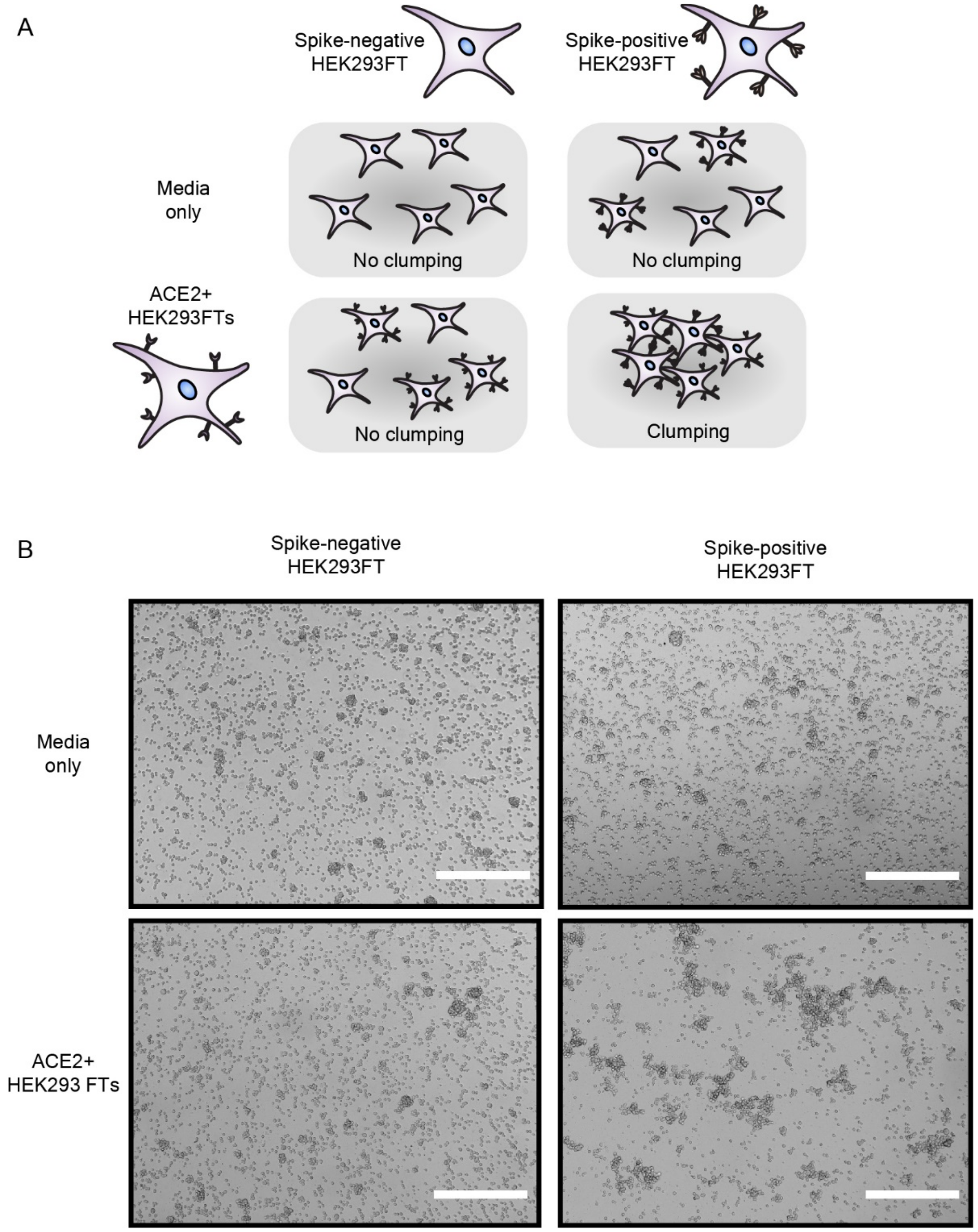


**Supplementary Figure 2.** Cells expressing Spike and ACE2 bind and clump. A) Cartoon depicting the cell binding assay. Cells expressing ACE2 or a media-only control sample are mixed with freshly plated Spike-positive or Spike-negative cells such that the total number of cells is equal for all conditions. Spike-positive and Spike-negative cells are a HEK293FT cell line engineered to inducibly express the Spike protein in the presence of doxycycline. Spike-positive cells in this assay have been cultured with doxycycline for greater than 24 h prior to assay, while Spike-negative cells are not cultured in doxycycline. If the binding interaction is specific, clumping is only expected for conditions where both ACE2 and Spike are present in trans. B) Representative microscopy images demonstrating cell binding as depicted in A. Scale bar is 500 μm.

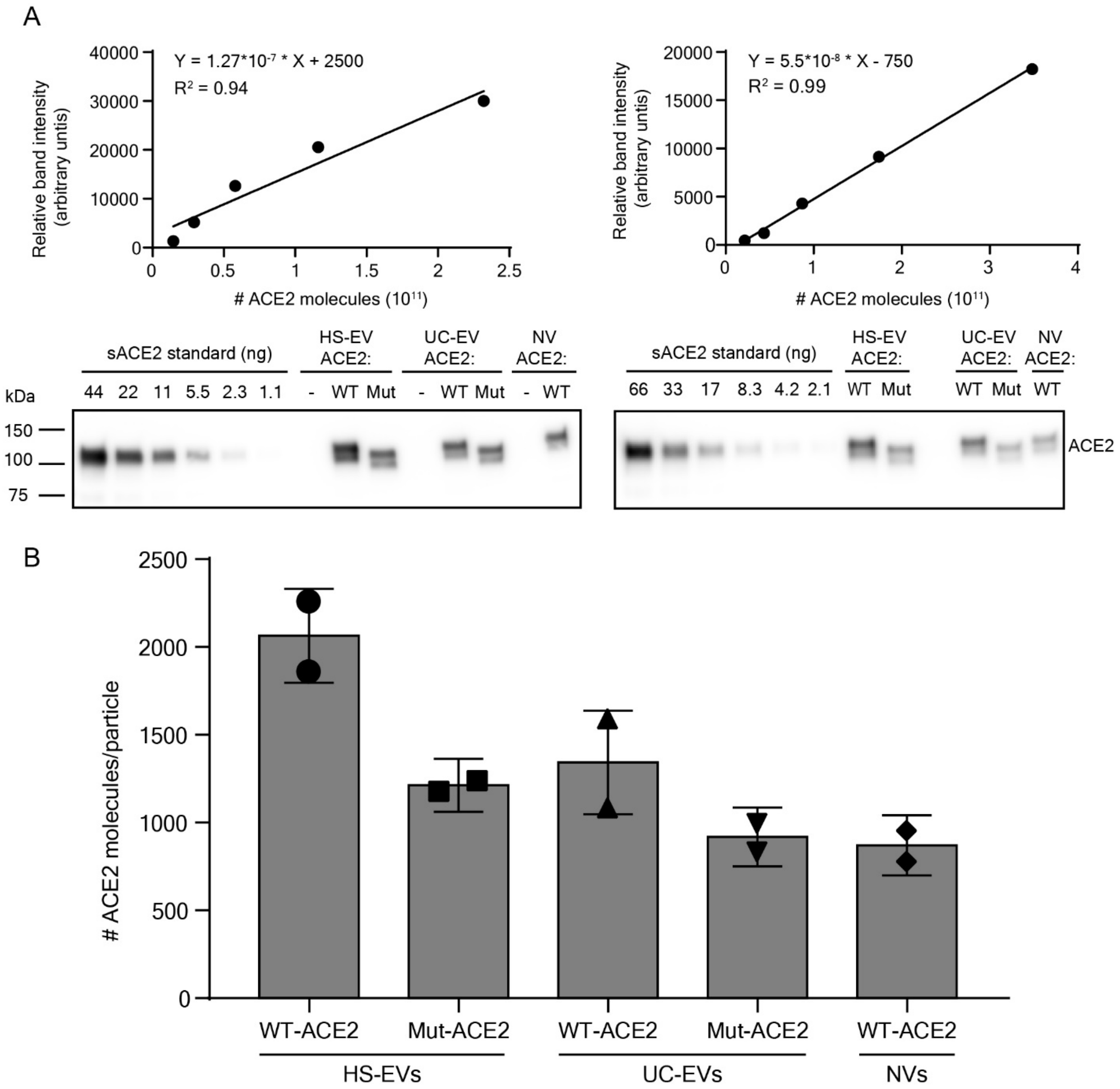


**Supplementary Figure 3.** Semi-quantitative western blots of the ACE2 content of vesicles. A) sACE2 was run on a western blot against an equal number of different vesicle subtypes. Calibration curves were generated from the intensity of sACE2 bands on the western blots. Equations for each curve are shown. Two independent replicates of this experiment were completed. B) ACE2 quantity per vesicle using the calibration curves derived in A. Data points represent a single value calculated from a single calibration curve; bar graphs represent the mean of the two individual calculated values. Error bars are the square root of the sum of the variances due to the standard error of the mean between the two data points and the propagated error from the individual measurements.

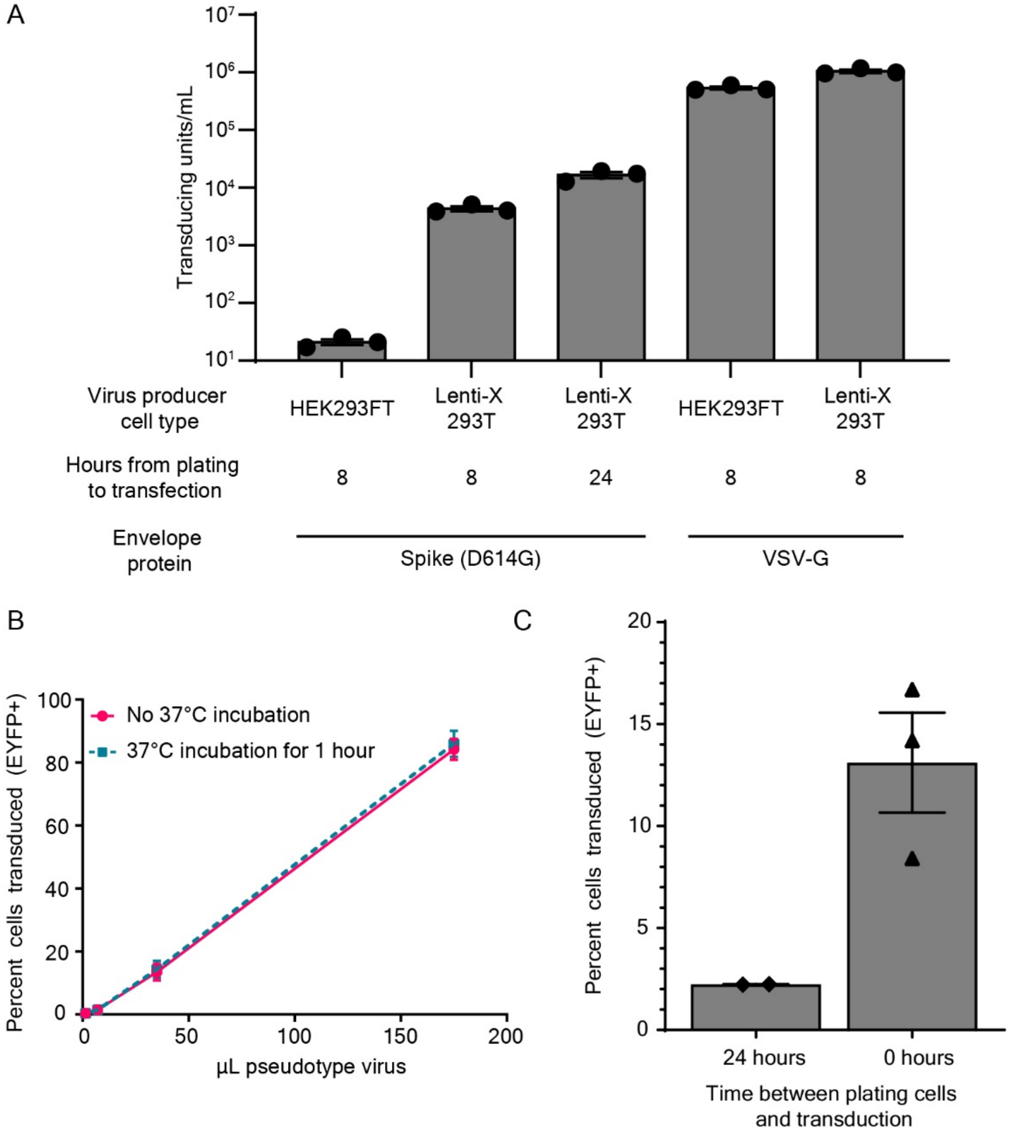


**Supplementary Figure 4.** Viral producer cell type and transduction conditions affect SARS-CoV-2 Spike-pseudotyped lentivirus titer. A) Functional viral titer for Spike-pseudotyped lentivirus (Spike-lenti) and VSV-G pseudotyped lentivirus generated from HEK293FT and HEK293T Lenti-X cells. Viral producing cells were plated at the time indicated before transfection of viral plasmids. Virus was titered on WT-ACE2 expressing HEK293FTs and determined using flow cytometry. Titer is presented as transducing units (TU) / mL of unconcentrated viral preparation.

Each symbol represents a biological replicate, the bar graph represents the mean of those replicates, and the error bars represent the standard error of the mean. B) Percent of cells transduced with different amounts of Spike-lenti that was either exposed to 37°C for 1 h or left on ice prior to transduction. Symbols represent the average from three biological replicates; error bars represent standard deviation. C) Percent of cells transduced for Spike-lenti for target cells plated at different times prior to inoculation; $4x10^3$ target cells were plated in both conditions and all conditions received the same volume of virus. ACE2-expressing HEK293FTs were more susceptible to transduction when plated at the time of transduction; effectively increasing viral titer. Spike-lenti in C) was harvested from HEK293FTs at 44 h after the media change; this virus not incubated for 1 h at 37°C or centrifuged at 500g for 1 min. Media was not changed after 16 h as in other experiments. Each symbol represents a biological replicate, the bar graph represents the mean of those replicates, and the error bars represent the standard error of the mean.

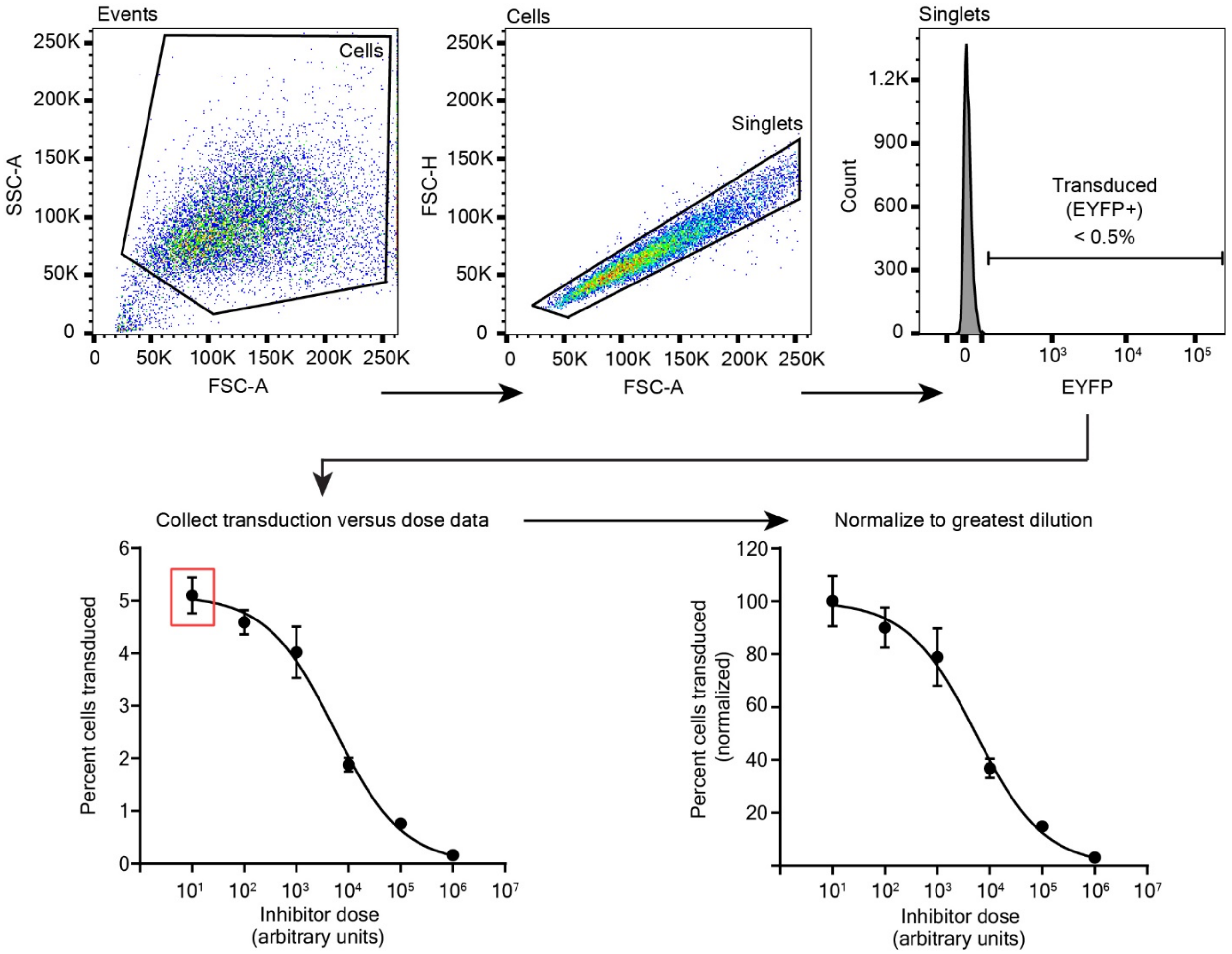


**Supplementary Figure 5.** Data analysis workflow for viral inhibition experiments. First, events recorded via analytical flow cytometry are gated for cells (via side scatter-area (SSC-A) vs forward scatter-area (FSC-A) discrimination) and then for single cells (via forward scatter-height (FSC-H) vs FSC-A discrimination). Transduced cells are then determined as cells that have higher EFYP fluorescence than 99.5% of single cells in a population not exposed to virus. Percent of cells transduced, as determined by the aforementioned gating, are then plotted against inhibitor dose. Percent cells transduced is then normalized to the percent transduction in the greatest dilution in a given treatment's series, here indicated with a red square.

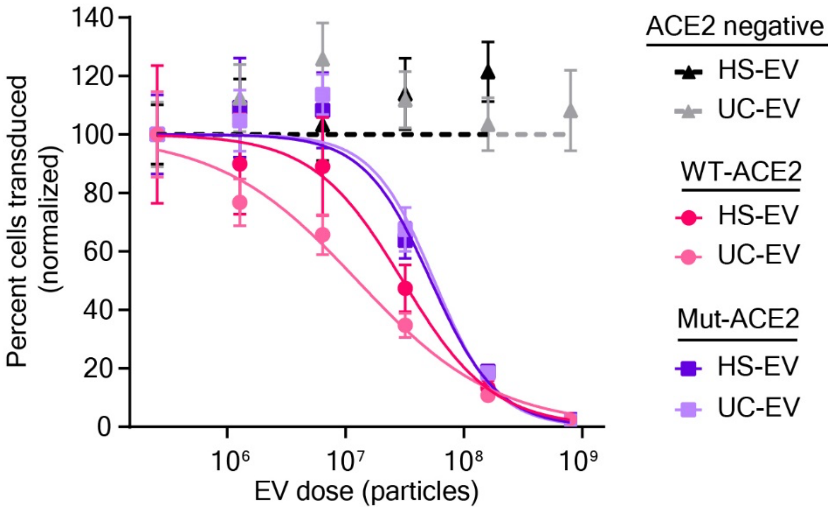


**Supplementary Figure 6.** Decoy EVs inhibit Spike-lenti. These data represent an independent, replicated experiment of **Figure 3D**. Dose-response curve demonstrating the relationship between decoy vesicle dose and normalized percentage of cells transduced by Spike-lenti. Curves are normalized to the percent of cells transduced at the lowest vesicle dose in a particular curve. Symbols repesent the mean of three biological replicates, and the error bars are standard error of the mean.

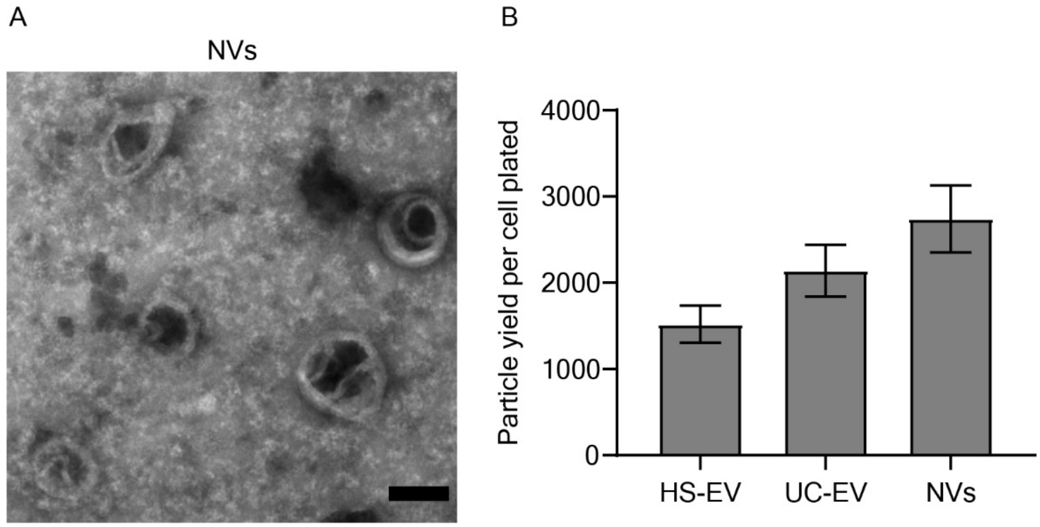


**Supplementary Figure 7.** NVs display unique morphology and provide similar particle yields to EVs. A) Transmission electron microscopy images depicting the presence of suspected multilamellar vesicles in a NV sample. Scale bar is 100 nm. B) Particle yield for NVs versus HS-EVs and UC-EVs normalized to the initial number of wild-type ACE2-engineered cells plated for the preparation. Error bars represent the standard deviation of a single vesicle preparation based on propagated error.

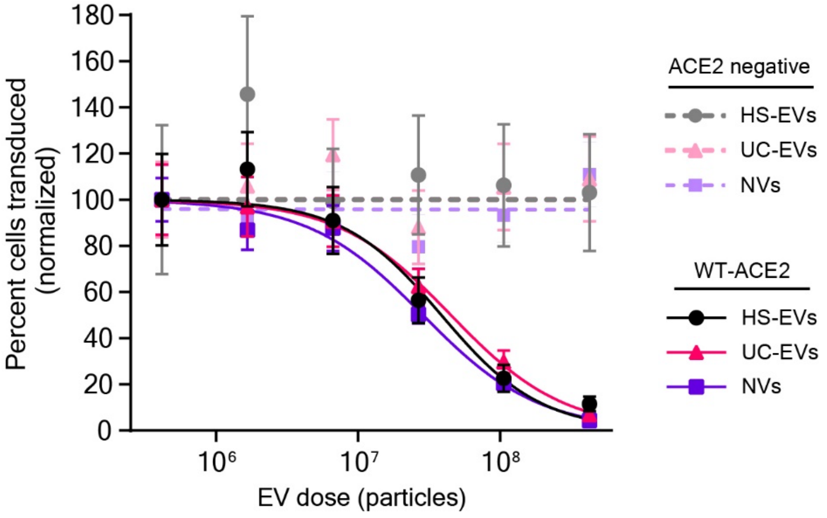


**Supplementary Figure 8.** NVs inhibit Spike-lenti. Independent, replicated experiment of **Figure 4F**. Dose-response curve demonstrating the relationship between decoy vesicle dose and normalized percentage of cells transduced by Spike-lenti. Curves are normalized to the percent of cells transduced at the lowest vesicle dose in a particular curve. Symbols repesent the mean of three biological replicates except for WT-ACE2 HS-EVs, where the third dilution in the series is the mean of two biological replicates. Error bars are standard error of the mean.

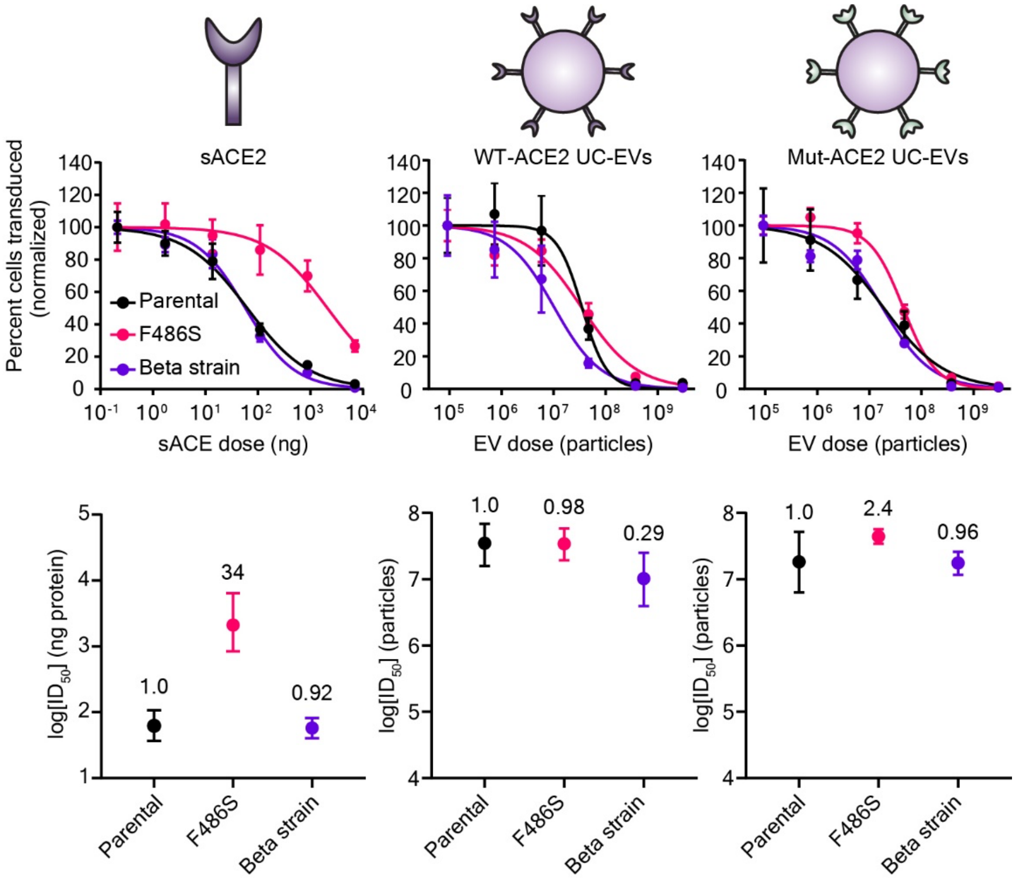


**Supplementary Figure 9.** Decoy vesicles maintain inhibition potency against Spike-lenti mutants. These data represent an independent, replicated experiment of **Figure 5B**. Dose-response curve demonstrating the relationship between decoy vesicle dose and normalized percentage of cells transduced by Spike-lenti. Curves are normalized to the percent of cells transduced at the lowest vesicle dose in a particular curve. Symbols repesent the mean of three biological replicates, and the error bars are standard error of the mean.

**Supplementary Table 1.** Calculated $ID_{50}$ and $IC_{50}$ parameters and errors for viral inhibition experiments investigating the effect of ACE2 mutant and EV subtype on Spike-lenti transduction corresponding to **Figure 3D**.

| | $ID_{50}$ (x $10^7$ Particles) | | $IC_{50}$ (x $10^7$ Particles) | |
|---|---|---|---|---|
| Sample | Best Fit | Uncertainty (95% C.I.) [-,+] | Best Fit | Uncertainty (95% C.I.) [-,+] |
| WT-ACE2 HS-EV | 2.8 | [0.7, 1] | 14 | [4, 5] |
| WT-ACE2 UC-EV | 3.8 | [0.9, 1] | 19 | [4, 6] |
| Mut-ACE2 HS-EV | 1.3 | [0.5, 0.7] | 6.5 | [2, 3] |
| Mut-ACE2 UC-EV | 2.2 | [0.4, 0.5] | 11 | [2, 2] |

**Supplementary Table 2.** Calculated $ID_{50}$ and $IC_{50}$ parameters and errors for viral inhibition experiments investigating the effect of ACE2 mutant and EV subtype on Spike-lenti transduction corresponding to **Figure S6**.

| | $ID_{50}$ (x $10^7$ Particles) | | $IC_{50}$ (x $10^7$ Particles) | |
|---|---|---|---|---|
| Sample | Best Fit | Uncertainty (95% C.I.) [-,+] | Best Fit | Uncertainty (95% C.I.) [-,+] |
| WT-ACE2 HS-EV | 3.0 | [2, 4] | 15 | [8, 20] |
| WT-ACE2 UC-EV | 1.3 | [0.5, 0.9] | 6.3 | [3, 5] |
| Mut-ACE2 HS-EV | 5.3 | [2, 4] | 26 | [10, 20] |
| Mut-ACE2 UC-EV | 5.6 | [2, 3] | 28 | [9, 10] |

**Supplementary Table 3.** Calculated $ID_{50}$ and $IC_{50}$ parameters and errors for viral inhibition experiments investigating the potency of wild-type ACE2 EVs versus cell-derived nanovesicles on Spike-lenti transduction corresponding to **Figure 4F**.

| | $ID_{50}$ (x $10^7$ Particles) | | $ID_{50}$ (x $10^7$ Particles) | |
|---|---|---|---|---|
| Sample | Best Fit | Uncertainty (95% C.I.) [-,+] | Best Fit | Uncertainty (95% C.I.) [-,+] |
| HS-EVs | 2.3 | [0.7, 1] | 11 | [3, 5] |
| UC-EVs | 3.6 | [1, 2] | 18 | [7, 10] |
| NV | 2.0 | [0.6, 0.8] | 10. | [3, 4] |

**Supplementary Table 4.** Calculated $ID_{50}$ and $IC_{50}$ parameters and errors for viral inhibition experiments investigating the potency of wild-type ACE2 EVs versus cell-derived nanovesicles on Spike-lenti transduction corresponding to **Figure S8**.

| | $ID_{50}$ (x $10^7$ Particles) | | $ID_{50}$ (x $10^7$ Particles) | |
|---|---|---|---|---|
| Sample | Best Fit | Uncertainty (95% C.I.) [-,+] | Best Fit | Uncertainty (95% C.I.) [-,+] |
| HS-EVs | 3.9 | [2, 4] | 19 | [10, 20] |
| UC-EVs | 4.5 | [2, 3] | 22 | [9, 20] |
| NV | 2.9 | [0.9, 1] | 14 | [5, 7] |

**Supplementary Table 5.** Calculated $ID_{50}$, $IC_{50}$, $ID_{50}$/TU and relative resistance parameters for viral inhibition experiments investigating the potency of UC-EVs and soluble ACE2 against Spike-lenti mutants corresponding to **Figure 5B**.

| Inhibitor | Virus | $ID_{50}$ (x $10^7$ particles or ng protein) | Relative resistance | $IC_{50}$ (x $10^7$ particles/mL or ng protein/mL) | $ID_{50}$/TU (particles/TU or ng protein/TU) | Relative resistance-TU normalized |
|---|---|---|---|---|---|---|
| sACE2 | Parental | 83 | 1.0 | 420 | 0.55 | 1.0 |
| | sACE2 Resistant | 4400 | 53 | 22000 | 46 | 84 |
| | Beta Strain RBD | 43 | 0.51 | 210 | 0.18 | 0.34 |
| WT-ACE2 UC-EVs | Parental | 2.0 | 1.0 | 9.9 | 200000 | 1.0 |
| | sACE2 Resistant | 1.5 | 0.76 | 7.5 | 130000 | 0.66 |
| | Beta Strain RBD | 0.32 | 0.16 | 1.6 | 13000 | 0.066 |
| Mut-ACE2 UC-EVs | Parental | 1.7 | 1.0 | 8.6 | 120000 | 1.0 |
| | sACE2 Resistant | 1.6 | 0.93 | 8.0 | 140000 | 1.2 |
| | Beta Strain RBD | 1.1 | 0.62 | 5.3 | 52000 | 0.42 |

**Supplementary Table 6.** Calculated $ID_{50}$, $IC_{50}$, and $ID_{50}$/TU parameters converted to numbers of molecules of ACE2 for viral inhibition experiments investigating the potency of UC-EVs and soluble ACE2 against Spike-lenti mutants corresponding to **Figure 5B**.

| Inhibitor | Virus | $ID_{50}$ (x $10^9$ molecules) | $IC_{50}$ (x $10^9$ molecules/mL) | $ID_{50}$/TU (x $10^9$ molecules/TU) |
|---|---|---|---|---|
| sACE2 | Parental | 440 | 2200 | 2.9 |
| | sACE2 Resistant | 23000 | 120000 | 240 |
| | Beta Strain RBD | 220 | 1100 | 0.96 |
| WT-ACE2 UC-EVs | Parental | 27 | 130 | 0.26 |
| | sACE2 Resistant | 20 | 100 | 0.17 |
| | Beta Strain RBD | 4.3 | 22 | 0.017 |
| Mut-ACE2 UC-EVs | Parental | 16 | 79 | 0.11 |
| | sACE2 Resistant | 15 | 73 | 0.13 |
| | Beta Strain RBD | 9.8 | 49 | 0.048 |

**Supplementary Table 7.** Calculated $ID_{50}$, $IC_{50}$, $ID_{50}$/TU and relative resistance parameters for viral inhibition experiments investigating the potency of UC-EVs and soluble ACE2 against Spike-lenti mutants corresponding to **Figure S9**.

| Inhibitor | Virus | $ID_{50}$ (x $10^7$ particles or ng protein) | Relative resistance | $IC_{50}$ (x $10^7$ particles/mL or ng protein/mL) | $ID_{50}$/TU (particles/TU) or (ng protein/TU) | Relative resistance-TU normalized |
|---|---|---|---|---|---|---|
| sACE2 | Parental | 62 | 1.0 | 310 | 0.30 | 1.0 |
| | sACE2 Resistant | 2100 | 34 | 11000 | 15 | 50 |
| | Beta Strain RBD | 58 | 0.92 | 290 | 0.42 | 1.4 |
| WT-ACE2 UC-EVs | Parental | 3.5 | 1.0 | 18 | 290000 | 1.0 |
| | sACE2 Resistant | 3.4 | 0.98 | 17 | 230000 | 0.79 |
| | Beta Strain RBD | 1.0 | 0.29 | 5.1 | 66000 | 0.23 |
| Mut-ACE2 UC-EVs | Parental | 1.8 | 1.0 | 9.2 | 120000 | 1.0 |
| | sACE2 Resistant | 4.4 | 2.4 | 22 | 340000 | 2.9 |
| | Beta Strain RBD | 1.8 | 0.96 | 8.8 | 150000 | 1.3 |

**Supplementary Table 8.** Calculated $ID_{50}$, $IC_{50}$, and $ID_{50}$/TU parameters converted to numbers of molecules of ACE2 for viral inhibition experiments investigating the potency of UC-EVs and soluble ACE2 against Spike-lenti mutants corresponding to **Figure S9**.

| Inhibitor | Virus | $ID_{50}$ (x $10^9$ molecules) | $IC_{50}$ (x $10^9$ molecules/mL) | $ID_{50}$/TU (x $10^9$ molecules/TU) |
|---|---|---|---|---|
| sACE2 | Parental | 330 | 1600 | 1.6 |
| | sACE2 Resistant | 11000 | 55000 | 78 |
| | Beta Strain RBD | 300 | 1500 | 2.2 |
| WT-ACE2 UC-EVs | Parental | 47 | 230 | 0.38 |
| | sACE2 Resistant | 46 | 230 | 0.3 |
| | Beta Strain RBD | 14 | 69 | 0.088 |
| Mut-ACE2 UC-EVs | Parental | 17 | 84 | 0.11 |
| | sACE2 Resistant | 41 | 200 | 0.31 |
| | Beta Strain RBD | 16 | 81 | 0.14 |

**Supplementary Table 9.** Calculated $ID_{50}$, $IC_{50}$, $ID_{50}$/TU and relative resistance parameters for viral inhibition experiments investigating the potency of WT-ACE2 UC-EVs against Spike-lenti mutants corresponding to **Figure 6**.

| Virus | $ID_{50}$ (x $10^7$ particles) | Relative resistance | $IC_{50}$ (x $10^7$ particles/mL) | $ID_{50}$/TU (particles/TU) | Relative resistance-TU normalized |
|---|---|---|---|---|---|
| Parental | 3.1 | 1 | 15 | 81000 | 1 |
| Delta | 2.1 | 0.7 | 11 | 79000 | 0.98 |
| Delta-plus | 1.4 | 0.47 | 7.1 | 70000 | 0.87 |
| Lambda | 2.4 | 1 | 12 | 87000 | 1 |

**Supplementary Table 10**. Western blot antibodies used in this study and associated sample preparation considerations.

| Antibody target | Supplier (#) | Boil Temp/Time | Antibody dilution | Reducing/Non-reducing Laemmli | Animal of origin |
|---|---|---|---|---|---|
| HA-tag | Cell Signaling Technology (C29F4) | 70°C / 10 min | 1:1000 | Reducing | Rabbit |
| ACE2 (C-term) | Abcam (Ab15348) | 70°C / 10 min | 1:1000 | Reducing | Rabbit |
| ACE2 (N-term)* | R&D Systems (MAB933) | 95°C / 5 min | 1:1000 | Reducing | Mouse |
| CD9 | Santa Cruz (sc-13118) | 95°C / 10 min | 1:500 | Reducing | Mouse |
| CD81 | Santa Cruz (sc-23962) | 95°C / 10 min | 1:500 | Non-reducing | Mouse |
| Alix | Abcam (Ab117600) | 95°C / 10 min | 1:500 | Reducing | Mouse |
| Calnexin | Abcam (Ab22595) | 70°C / 10 min | 1:1000 | Reducing | Rabbit |
| Rabbit | Invitrogen (32460) | Not applicable | 1:3000 | Not applicable | Goat |
| Mouse | Cell Signaling Technology (7076) | Not applicable | 1:3000 | Not applicable | Horse |

* Resuspended in PBS at recommended concentration (0.5 mg/mL)
